# WEGS: a cost-effective sequencing method for genetic studies combining high-depth whole exome and low-depth whole genome

**DOI:** 10.1101/2023.04.27.538531

**Authors:** Claude Bhérer, Robert Eveleigh, Katerina Trajanoska, Janick St-Cyr, Antoine Paccard, Praveen Nadukkalam Ravindran, Elizabeth Caron, Nimara Bader Asbah, Clare Wei, Iris Baumgartner, Marc Schindewolf, Yvonne Döring, Danielle Perley, François Lefebvre, Pierre Lepage, Mathieu Bourgey, Guillaume Bourque, Jiannis Ragoussis, Vincent Mooser, Daniel Taliun

## Abstract

Whole genome sequencing (WGS) at high-depth (30X) allows the accurate discovery of variants in the coding and non-coding DNA regions and helps elucidate the genetic underpinnings of human health and diseases. Yet, due to the prohibitive cost of high-depth WGS, most large-scale genetic association studies use genotyping arrays or high-depth whole exome sequencing (WES). Here we propose a novel, cost-effective method, which we call “Whole Exome Genome Sequencing” (WEGS), that combines low-depth WGS and high-depth WES with up to 8 samples pooled and sequenced simultaneously (multiplexed). We experimentally assess the performance of WEGS with four different depth of coverage and sample multiplexing configurations. We show that the optimal WEGS configurations are 1.7-2.0 times cheaper than standard WES (no-plexing), 1.8-2.1 times cheaper than high-depth WGS, reach similar recall and precision rates in detecting coding variants as WES, and capture more population-specific variants in the rest of the genome that are difficult to recover when using genotype imputation methods. We apply WEGS to 862 patients with peripheral artery disease and show that it directly assesses more known disease-associated variants than a typical genotyping array and thousands of non-imputable variants per disease-associated locus.

## 1 Introduction

Accurate assessment of DNA sequence variation enables insights into the genetic basis of diseases and other traits. Whole genome sequencing (WGS) at high-depth of coverage (30X and above) using next generation sequencing technologies is the current gold standard method for the accurate discovery of single nucleotide variants (SNVs) and short insertions/deletions (InDels) genome-wide^1, 2^. Sequencing offers several advantages over array-based genotyping, notably that variant positions are not fixed, which allows the discovery of novel population-specific variants. Yet, despite the decreasing costs of high-depth WGS, sequencing a large number of samples remains expensive. So far, the use of whole exome sequencing (WES) has dominated large-scale sequencing studies such as gnomAD^3^ and UK Biobank^4^, but WES is limited to coding regions. As a result, there is still a need for more cost-effective solutions to capture both coding and non-coding variation.

The array-based genotyping coupled with genotype imputation at untyped genomic positions from public haplotype reference panels^2, 5, 6^ is a popular, cost-effective strategy for increasing statistical power and genomic coverage in current genome-wide association studies (GWAS)^7^. The largest TOPMed haplotype reference panel allows for the imputation of variants down to minor allele frequencies (MAF) of ∼0.002–0.003% (imputation quality r^2>0.3) in individuals of European and African ancestries^6^. However, rare variant imputation with TOPMed still has much lower accuracy than common variant imputation, especially in non-European or non-African ancestry groups^6^. At the same time, the advantage of local sequencing-based imputation reference panels was demonstrated for multiple populations, such as the Estonian^8^, Finnish^9^ and Sardinian^10^.

Several cost-effective sequencing-and-imputation strategies have been described to improve genomic coverage while allowing better assessment of population-specific variants. Those include (a) WGS in a subset of study participants (at a depth ranging from 5X to 30X) to create a customized reference panel^7^ for imputation of the remaining participants who were genotyped using genotyping arrays and (b) ultra-low depth WGS (depth of coverage (DP) down to 0.1X-0.5X) or (c) low-depth (1X-4X) WGS in all study participants followed by imputation using public reference panels^11–14^. While ultra-low depth WGS can be performed at the same cost as array-based genotyping^11^, it has also been suggested that ultra-low depth and low-depth sequencing plus imputation are good alternative technologies to imputed genotyping arrays by doubling the number of true association signals discovered^14^ and improving the accuracy of polygenic risk prediction models^12, 13^. The latter models have also benefited from the inclusion of rare coding variants in their prediction algorithms^15–17^. However, recent work suggested that array-based imputation strategies may miss approximately half of the rare coding variants with MAF<0.05% detected by WES^2^. Although cheaper than WGS, WES is still a more expensive option than imputation-based strategies, and it ignores the majority of non-coding regions of the genome. Assessment of genetic variation in non-coding regions, which contains the vast majority of genetic variants^2^ and a majority (84%) of GWAS association signals^18^, is critical for many genetic analyses, notably understanding regulatory genetic variation.

Here, we propose a novel sequencing method, which we call Whole Exome Genome Sequencing (WEGS), that combines low-depth WGS (2-5X) and high-depth WES (100X) with up to 8 samples pooled and sequenced simultaneously (multiplexed) to reduce reagents costs^19^. We experimentally demonstrate that WEGS, while being 1.7-2.0 times cheaper than standard high-depth WES (100X) due to multiplexing and 1.8-2.1 times cheaper than 30X WGS, maintains similar precision and recall rates in the discovery of rare coding variants and allows assessment of population-specific variants in the rest of the genome. We demonstrate the scalability and utility of WEGS by applying it to 862 patients with peripheral artery disease (PAD).

## 2 Methods

### 2.1. DNA samples for benchmarking experiments

To benchmark our new method, we used DNA samples derived from cell lines obtained from the US National Institute of Standards and Technology (NIST) RM 8392, a family trio of Ashkenazi Jewish origin including a son (HG002), father (HG003) and mother (HG004), consented by the Personal Genome Project (PGP)^20^. These DNA samples were developed for the Genome in a Bottle (GIAB) Consortium to generate reference datasets for benchmarking genomic analyses^21^, and have broad, open consent for all research uses under the terms of the PGP.

### 2.2. Benchmarking experimental study design

To assess the relative performance of different WEGS protocols, we used DNA samples from the Ashkenazi trio to perform a series of WES and low-depth WGS sequencing experiments. For WES, we performed experiments without and with multiplexing of 4 and 8 samples (no-plexing, 4-plexing and 8-plexing, correspondingly). For each sample in the family trio, we performed library preparation and sequencing to a target DP of 100X in triplicate for the 1-plex and 4-plex WES experiments, and in duplicate for the 8-plex experiment, for a total of 37 samples (Figure 1). For WGS, using pre-capture libraries prepared for WES, we sequenced the trio samples to a target DP of 5X on 2 separate lanes. This allowed us to use a single lane to obtain a target DP of 2.5X. This gave us the possibility to evaluate four WEGS combinations: WEGS_4P,2X_, WEGS_4P,5X_, WEGS_8P,2X_, and WEGS_8P,5X_, where 4P and 8P denote 4-and 8-plexing, respectively, and 2X and 5X correspond to target DP of WGS.

**Figure 1.**
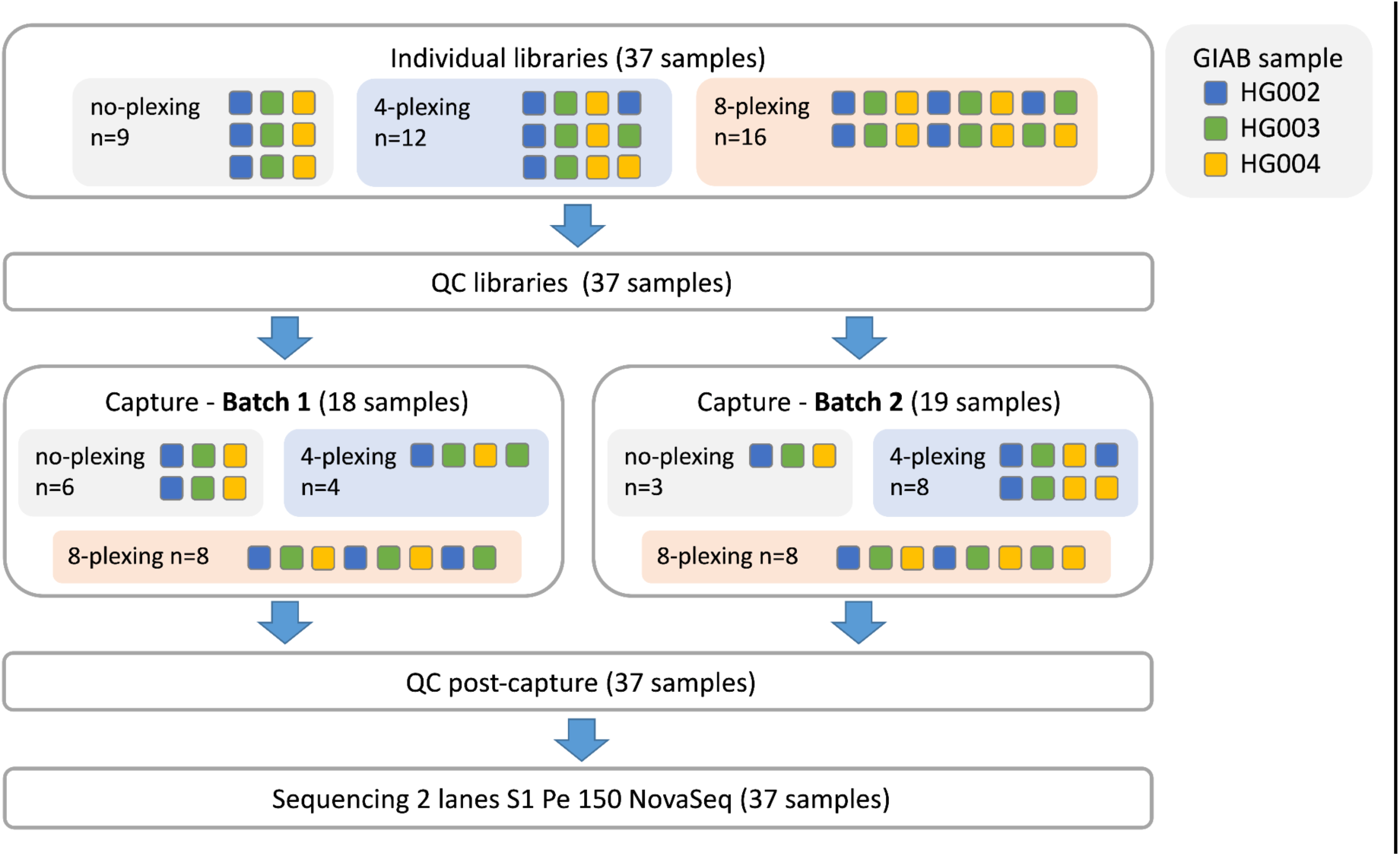
WES experimental design overview. DNA samples from a GIAB family trio (HG002, HG003, HG004) were used to perform WES experiments without and with multiplexing of 4 and 8 samples (no-plexing, 4-plexing and 8-plexing, correspondingly). For each sample in the family trio, we performed library preparation and sequencing to a target coverage of 100X in triplicate for the no-plexing and 4-plexing WES experiments, and in duplicate for the 8-plexing experiment, for a total of 37 samples. Sequencing library QC was performed before and after exome capture. Sequencing was performed using the Illumina NovaSeq S1 platform.

### 2.3. Sequence data production

WES and WGS sequencing was performed at the McGill Genome Centre in October 2021. Processing included sample quality control (QC) using a QUBIT 1X DSDNA HS ASSAY KT from Life Technologies Inc .to measure DNA concentration quality. An aliquot of 200 ng input in 50 ul total was used to perform DNA fragmentation (shearing) with Covaris LE220 (Covaris Inc.) method to a target of 300 bp fragments. Sample library preparation was carried out using Agilent SureSelect XT HS2. Subsequent captures were performed using Agilent SureSelect XT HS2 V7 capture panel with different plexing strategies: 4-plex (12 samples) and 8-plex (16 samples) (Figure 1). Unique dual sample indexing barcodes (2×8bp) were added to multiplexed samples during library preparation. Library QC was performed before and after capture in 2 steps: quantification using qPCR (Kapa Biosystems, part #KK4602) and QC using LabChip GX Touch HT Nucleic Acid Analyzer. Exome captures were performed in 2 batches using Agilent SureSelect Human All Exon V7 capture for a total 48.2-Mb target. Sequencing was performed on 2 lanes of the Illumina NovaSeq platform using S1 flowcells and 150-bp paired-end reads to a target coverage of 100X. Sample pre-capture libraries were used to perform WGS sequencing to a target coverage of 5X in 2 separate lanes on the Illumina NovaSeq platform using S1 flowcells to 150-bp paired-end reads.

### 2.4. Data processing and variant calling

As defined by Genome Analysis Tool Kit^22^ (GATK v4.2.0.0) best practice recommendations, preprocessed reads trimmed by the removal of adapters and low quality bases, were aligned to the decoy version of GRCh37 human genome build (hs37d5) using bwa-mem^23^ (v0.7.17) (Supplementary Figure 1). Mapped reads were further refined using GATK InDel realignment^22^ (v3.8) to improve the mapping of reads near InDels, marking of duplicated reads using GATK mark duplicates, and improve base quality scores using Base Quality Score Recalibration (BQSR). For WEGS processing, WGS and WES were analyzed by applying the above methods but using different trimming and mark duplication procedures to take advantage of the UMIs present in the WES data. The trimmer and locatIT programs from Agilent’s AGeNT tool set (v2.0.5) were used to first identify and remove the adaptor sequences, extract the molecular barcodes (MBC), and then merge duplicated reads by leveraging the MBC information embedded in the aligned BAM file. WGS data were processed using the read trimmer skewer^24^ (v0.2.2), and duplicated reads were assessed using GATK mark duplicates. Variant calling for all the experiments was performed using the GATK’s HaplotypeCaller.

### 2.5. Benchmark variant calls and regions

Benchmark (or “high-confidence”) variant calls for SNVs and short InDels from GIAB Consortium for each sample in the Ashkenazi family trio were obtained for build GRCh37 (v.4.2.1)^25^ at URL: https://ftp-trace.ncbi.nlm.nih.gov/ReferenceSamples/giab/release/. We used Illumina hap.py benchmarking tool (version v.0.3.10) to compare our study variant calls and imputed variants to GIAB “high-confidence” variant calls in previously described “high-confidence regions”^26, 27^. Variant calling recall rate was estimated as the total number of true positive variant calls divided by the total number of variant calls, and precision as the total number of true positive variant calls over the sum of true positive and false positive variant calls. We used imputed best-guess genotypes when estimating recall and precision rates for imputed variants against GIAB “high-confidence” variant calls. Before benchmarking, we did not apply filters on our variant calls (using variant calling annotations) or imputed genotypes (using imputation quality scores) to limit the contribution of other factors when interpreting differences between methods.

### 2.6. High-depth WGS data

To generate 30X WGS data for the Ashkenazi Jewish trio (HG002, HG003, and HG004), we downloaded 300X WGS data from GIAB produced using Illumina HiSeq 2500 in Rapid mode (v1) (PCR-free, pair-end, mean read length 2 x 148bp). The reads were aligned to the GRCh37 genome build using Novoalign version 3.02.07. Then, we randomly subset 10% of the reads using the samtools^28^ tool to reach 30X coverage on average. For each individual, we generated five 30X WGS datasets using different random seeds. Then, we performed variant calling using GATK v4.2 in the same way as for other experiments.

### 2.7. Tests for statistical significance

We used Wilcoxon rank-sum test to test for statistical significance (1) of differences between two library preparation batches and (2) of variant recall and precision rates between no-plexing WES and WEGS. We used Wilcoxon signed-rank test when comparing the same WES experiments (1) before and after UMI-aware read deduplication and (2) before and after adding WGS reads. We used a one-sided version of the tests depending on the means of two samples, i.e. the alternative hypothesis was that the distribution underlying the sample with a larger mean is stochastically greater than the distribution underlying the sample with a smaller mean. We used the implementation of both tests available in SciPy^29^. To assess the correlation strength between levels of multiplexing and different sequence data metrics, we used the Pearson correlation coefficient and the corresponding *P*-values for the two-sided alternative hypothesis that the correlation is non-zero implemented in SciPy^29^. We used *P*-value<0.05 for the statistical significance threshold.

### 2.8. Genotype imputation using genotyping arrays

To mimic genotyping array data for HG002, HG003, and HG004 samples, we subset 654,013 GRCh37 positions on the Infinium Global Screening Array 24 v3 (https://support.illumina.com/array/array_kits/infinium-global-screening-array/downloads.html) from the corresponding GIAB’s WGS data. Then, we imputed each sample individually using the multi-ethnic TOPMed reference panel (N=97,256) available at NHLBI TOPMed Imputation Server. In addition to genotype imputation, the server lifted positions from GRCh37 to GRCh38 genome build and performed reference-based statistical phasing. The imputed genotypes were on the GRCh38 genome build. To compare them to WEGS, we used the GATK LiftoverVcf tool^30^ to lift imputed positions back to GRCh37. We annotated the data after imputation with alternate allele frequencies (AF) in the Ashkenazi Jewish (ASJ) population from gnomAD v3.1.1^3^ and overall AF in the BRAVO variant browser, which includes all individuals in the TOPMed reference panel. For both databases, we lifted the GRCh38 positions to GRCh37 using GATK LiftoverVcf. We used only those variants which passed all quality filters described by gnomAD and TOPMed, correspondingly. When comparing AF distributions in ASJ vs BRAVO, we restricted our analyses to non-monomorphic genetic variants where at least 1,000 ASJ individuals were sequenced.

### 2.9. Genotype imputation using WEGS and local reference panel

We used the GLIMPSE method^31^ to impute variants from the local reference panel using sequencing reads in WEGS. To build our local reference panel, we used genotypes from the 1000 Genomes Project (1000G)^6^ and Human Genome Diversity Project (HGDP)^32^ (N=4,150) from gnomAD v3^3^. We kept only variants which passed all quality filters defined by gnomAD v3, were missing in <1% of individuals, and for which alternate allele count was ≥2. We phased the genotypes using statistical phasing implemented in SHAPEIT4^33^ and lifted positions of phased genotypes from GRCh38 to GRCh37 genome build using the GATK LiftoverVcf tool. We merged the GLIMPSE-imputed variants with variants directly genotyped from WEGS by GATK. We kept the genotyped version when a variant was imputed and genotyped at the same time (i.e. had the same position and alleles).

### 3.0. WEGS application

A total of 862 patients diagnosed with PAD were recruited and consented between April 2, 2017, and September 21, 2021, in the Division of Angiology, at the Insel University Hospital of Bern, Switzerland. Recruited patients had whole blood samples collected and stored in the Liquid Biobank Bern (LBB). We applied the above WEGS method to each sample using WGS at an average depth close to 5X and WES at 100X. Exomes were captured in 8-plex using the Agilent SureSelect All Exons Human V7 capture. The exome and whole genome libraries were sequenced on MGI T7 sequencers. All sequence reads were mapped to build GRCh38. We followed GATK best practices pipelines for jointly calling SNVs and InDels. We used only those variants which passed all variant filters after GATK’s VQSR and had less than 1% missing genotypes.

## 3 Results

### 3.1. Sample multiplexing lowers depth of coverage due to duplicate reads

Sample multiplexing allows multiple samples to be pooled and sequenced simultaneously, resulting in lower per-sample sequencing costs^19^. However, multiplexing may also increase the number of false positive variant calls^34^. To assess sequencing quality, we first compared the DP and variant calling when using WES without and with multiplexing of 4 and 8 samples (no-plexing, 4-plexing and 8-plexing, correspondingly). For this, we generated 37 exome sequences at 100X WES and different levels of multiplexing using DNA from Ashkenazi trio samples (Figure 1, see Methods).

We observed a strong negative correlation (Pearson’s *r* = -0.69, *P*-value = 2.31⨉10^-6^) between the average DP in targeted exome regions and the number of multiplexed samples (Figure 2). The median values of average DP across individual exomes dropped from 121.8 in no-plexing experiments to 98.6 and 82.6 in 4-plexing and 8-plexing, respectively. The average DP ratio between no-plexing and 4-and 8-plexing experiments was similar in all targeted regions across the exome - showing no evidence that differences in average DP were non-uniform or affected only a subset of targeted regions (Supplementary Figure 2). When stratifying by library preparation batch, we observed statistically significant differences (*P*-value = 0.048) in average DP between two batches only in experiments without multiplexing (Supplementary Figure 3A). Nevertheless, these differences did not influence the overall trend - the strong negative correlations between the number of multiplexed samples and DP remained in both library preparation batches (Supplementary Figure 3B-C).

**Figure 2.**
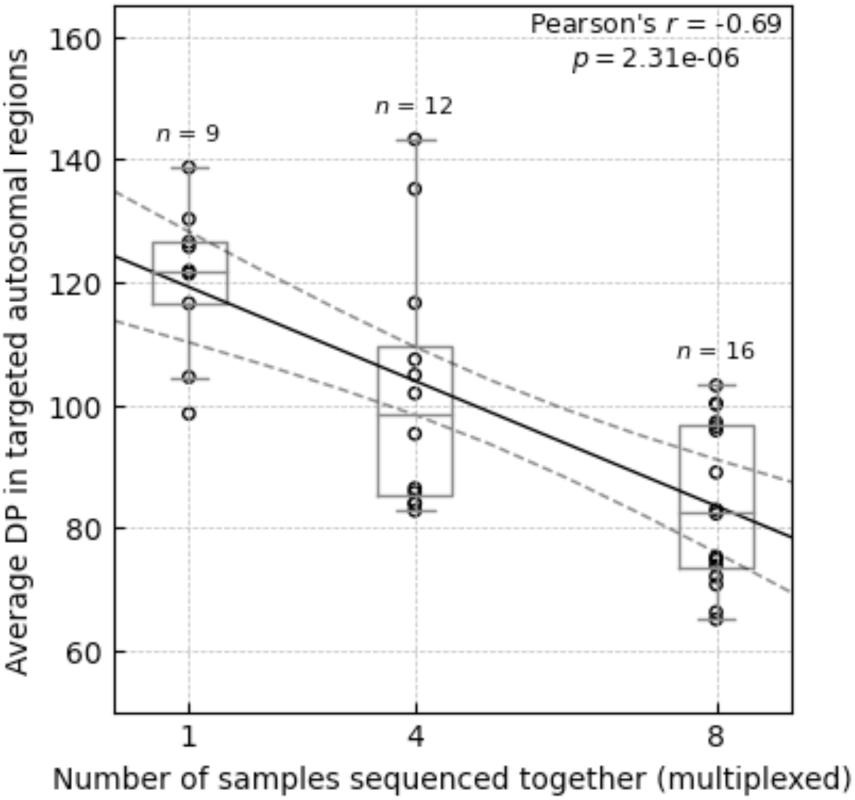
Average depths of coverage across all targeted regions in autosomal chromosomes in WES experiments without and with multiplexing. The average depth of coverage (DP) was computed across target regions in Agilent V7 capture using paired mapped reads and counting only base-pairs with minimal Phred-scaled mapping and base qualities of 20. The solid black line corresponds to the linear regression line, and the dashed black lines correspond to a 95% confidence interval. The box bounds the IQR, and Tukey-style whiskers extend to a maximum of 1.5 ⨉ IQR beyond the box. The horizontal line within the box indicates the median value. Open circles are data points corresponding to the average DP across individual exome.

To better understand the cause of lower average DP in multiplex sequencing, we assessed the total number of paired reads, the number of reads flagged as PCR or optical duplicates, the number of unmapped reads, and the average base qualities in reads. There was no correlation (Pearson’s *r* = -0.08, *P*-value = 0.631) between the total number of paired reads and the number of samples pooled together for sequencing (Supplementary Figure 4A). However, there was a strong positive correlation (Pearson’s *r* = 0.92, *P*-value = 2.13 ⨉ 10^-15^) between the percent of reads flagged as PCR or optical duplicates and degrees of multiplexing (Supplementary Figure 4B). Compared to the multiplexing-free sequencing experiments, the 4-plexing and 8-plexing experiments showed a 1.7-fold (18.4% vs 31.2%) and 2.3-fold (18.4% vs 43.0%) increase in the median percent of duplicated reads, respectively. The data also suggested a weak, non-statistically significant correlation (Pearson’s *r* = 0.32, *P*-value = 0.06) between the percent of unmapped reads and degrees of multiplexing (Supplementary Figure 4C). Also, the percent of unmapped reads did not exceed 0.11 percent of the total number of paired reads and, thus, did not contribute much to the differences in average DP. There was a moderate correlation (Pearson’s *r* = -0.52, *P*-value=1.09 ⨉ 10^-3^) between the average base qualities and the degree of multiplexing (Supplementary Figure 4C). However, when stratified by the library preparation batch and in contrast to the other metrics mentioned above, the first batch did not show the same correlation pattern (Supplementary Figure 5-8), suggesting that other factors may affect the base qualities. We conclude that the main contributor to the lower average DP in sample multiplexing experiments compared to the experiments without sample multiplexing is the percent of reads flagged as PCR or optical duplicates.

### 3.2. UMI does not recover losses in the depth of coverage

UMI - a unique barcode appended to each DNA fragment before the PCR - helps to distinguish the truly duplicated fragments originating from the same molecule from the very similar fragments originating from a different molecule^35, 36^. In addition, UMI-aware software tools for duplicate read removal help to identify and remove sequencing errors by grouping reads with the same UMI and creating a consensus read^37^. We applied the duplex UMI method in our sequencing experiments and evaluated the utility of LocatIt and UmiAwareMarkDuplicatesWithMateCigar (GATK+UMI) UMI-aware read deduplication tools with multiplexing. LocatIt reduced the average DP in experiments without and with multiplexing, compared to the UMI agnostic deduplication approach, while UMI+GATK increased the average DP (Supplementary Figure 9). We explain the different effects on average DP by the difference in strategies between these two tools. For example, in 8-plexing experiments, the GATK+UMI reduced the percent of duplicated reads on average by 0.4 (SE = 0.01), while LocatIt reduced it by 1.56 (SE = 0.03) (Supplementary Table 1). However, LocatIt, on average, marked an additional 4.38% of reads as QC failed, which included reads with low base qualities in their UMIs and single consensus read pairs without complementary pairs. This additional filtering in LocatIt resulted in lower average DP, fewer unmapped reads, and higher average base qualities. In summary, the UMI-aware read deduplication showed that the vast majority of the duplicated reads in multiplexing experiments are truly PCR/optical duplicates. UMI-aware deduplication didn’t help recover the loss in average DP in multiplexing experiments back to the levels of no-plexing experiments.

### 3.3. Sample multiplexing decreases variant recall rates

We observed moderate-to-strong negative correlations between the number of samples sequenced together and the recall rates for SNVs (Pearson’s *r* = -0.60, P-value =7.79⨉10^-5^) and InDels (Pearson’s *r* = -0.48, *P*-value=2.85⨉10^-3^) (Supplementary Figures 10A and 10C). The average recall rates dropped from 0.983 (SE = 0.0004) and 0.939 (SE = 0.003) in no-plexing experiments to 0.980 (SE = 0.0004) and 0.926 (SE = 0.003) in 8-plexing experiments for SNVs and InDels, respectively (Supplementary Table 2). In many instances, the recall rates were lower in the second library preparation batch, and some of these differences were statistically significant (Supplementary Figures 11A and 11D). Despite these differences, the statistically significant negative correlations between variant recall rates and the number of multiplexed samples were present in both library preparation batches (Supplementary Figures 11B-C and 11E-F).

We also observed a drop in precision for both variant types with the increased number of multiplexed samples, but unlike recall, the negative correlations were weaker and not statistically significant (Supplementary Figures 10B and 10D, Supplementary Table 2). The precision rates were similar between the library preparation batches (Supplementary Figures 12A and 12D), and they also did not show statistically significant correlations with the number of multiplexed samples when stratified by batch (Supplementary Figures 12B-C and 12E-F). Only in the first batch we saw a weak positive correlation (Pearson’s r = 0.27, *P*-value = 0.26) between precision and the number of multiplexed samples (Supplementary Figure 12B).

We looked into the number of true positive (TP), false positive (FP), and false negative (FN) variant calls to explain the statistically significant decrease in recall rates. We found the strongest correlation in the degree of multiplexing and the number of FN calls (Pearson’s *r* = 0.60, *P*-value=8.46⨉10^-5^ in SNVs and Pearson’s *r* = 0.44, *P*-value=6.34⨉10^-3^ in InDels), representing the true variants that are not detected (Supplementary Figure 13). For example, the average number of undetected true SNVs increased from 384 (SE = 8) in single-sample sequencing experiments to 446 (SE = 9) in 8-plexing experiments (Supplementary Table 2). On average, 65 (SE = 6) true SNVs missed in 8-plexing experiments were correctly identified across all no-plexing experiments for the corresponding sample, and 61 (SE = 6) of those had a higher depth of coverage in no-plexing experiments than in 8-plexing experiments (Supplementary Table 3). We conclude that the main driver for the decrease in recall rates is the drop in average DP in multiplexing experiments, which leads to the increased number of missed true variants.

### 3.4. UMI improves variant calling insufficiently

We investigated how the recall and precision rates changed in SNV calling after applying the UMI-aware duplicate read removal. We wanted to test if a more accurate read deduplication could partially compensate for the loss of variant recall rates in multiplexing experiments. As previously, we considered two UMI-aware deduplication tools: LocatIt and UmiAwareMarkDuplicatesWithMateCigar (GATK+UMI).

We observed small but statistically significant drops in the recall rates when using LocatIt in all samples at all levels of plexing (Supplementary Figure 14A). For example, on average, the paired difference in the same sample in the 8-plexing experiment between two recall rates, one measured after LocatIt and another measured after the UMI agnostic approach, was only-0.0008 (SE=0.0001) (Supplementary Table 4). The paired differences between the recall rates were consistently negative in all samples in the 8-plexing experiment, and this relationship was statistically significant (*P*-value = 2⨉10^-3^) (Supplementary Figure 14A). However, there was no consistent and statistically significant change in the precision rates: precision slightly increased in some samples but dropped in others (Supplementary Figure 14B). For instance, while, on average, a paired difference in the 8-plexing experiment between precision rates increased by 0.0002 (SE=0.0002) (Supplementary Table 4), the paired differences between precision rates were negative in 5 out of 16 samples and did not support this average increase (*P*-value = 0.85) (Supplementary Figure 14B). The statistically significant decreases and increases were also in the total number of called SNVs and the number of missed true SNVs (i.e. FN calls), respectively (Supplementary Table 4). In samples in the 8-plexing experiments, the average paired difference between the total numbers of called SNVs was -20 (SE=4) and between the numbers of missed true SNVs was 17 (SE=2). Although the average paired difference between the numbers of FP calls was -2 (SE=3) and suggested a decrease in the numbers of FP calls when using LocatIt, this relationship was not statistically significant (*P*-value ≥ 0.05). The reduced number of called SNVs is consistent with our previous observation of reduced average DP when using LocatIt due to additional read filtering.

When using GATK+UMI, we observed slight but statistically significant improvements in the SNV recall rates for samples in multiplexing experiments (Supplementary Figure 14C). In samples in the 8-plexing experiments, the average paired difference between recall rates was 0.0003 (SE<0.0001) (Supplementary Table 4), and the increase in recall rates was observed in the majority of samples and supported the statistical significance of the relationship (*P*-value=3.1⨉10^-5^) (Supplementary Figure 14C). At the same time, there was also a slight statistically significant drop in the precision rates at all levels of plexing (Supplementary Figure 14D). In the same samples in the 8-plexing experiments, the average paired difference between precision rates was -0.0014 (SE=0.0001) (Supplementary Table 4), and the decrease was consistent across all samples leading to the statistically significant relationship (*P*-value=1.5⨉10^-5^) (Supplementary Table 4). The observed increase in the number of called SNVs (e.g. M=39 [SE=2] in 8-plexing) and the number of FP calls (e.g. M=33 [SE=2] in 8-plexing) with a much smaller decrease in the number of FN calls (e.g. M=-6 [SE=1] in 8-plexing) (Supplementary Table 4) can explain the increase in recall and decrease in precision rates. The increase in the number of called SNVs is consistent with our previous observation of increased DP when using GATK+UMI.

In summary, while UMI-aware read deduplication can improve SNV recall or precision rates depending on the approach, this improvement appears minimal in the present experiment. It does not allow to recover these rates back to levels similar to no-plexing experiments.

### 3.5. WEGS significantly improves variant calling in multiplexed samples

To compensate for the losses in variant recall rates when performing multiplexed WES, we introduced reads from low-depth WGS before variant calling. We called this approach WEGS. We evaluated four combinations in comparison to no-plexing WES: (1) 4-plexing WES and WGS at 2X average DP (WEGS_4P,2X_), (2) 4-plexing WES and WGS at 5X average DP (WEGS_4P,5X_), (3) 8-plexing WES and WGS at 2X average DP (WEGS_8P,2X_), and (4) 8-plexing WES and WGS at 5X average DP (WEGS_8P,5X_). In each combination, we looked at the paired difference in the same sample between two recall rates, one measured after adding reads from WGS and another before.

Additional reads from 2X and 5X WGS improved variant recall rates in all multiplexing experiments, and the differences were statistically significant (*P*-value<0.05) (Supplementary Figures 15 and 16). For instance, the average paired difference in SNV recall rates in WEGS_8P,2X_ was 0.0031 (SE=0.0002) (Supplementary Table 5). This paired difference in recall rates was positive across all samples and, thus, supported the statistical significance of the observed increase in recall rates (*P*-value=1.5⨉10^-5^) (Supplementary Figure 15A). The total number of discovered SNVs increased on average by 76 (SE=6), of which 70 (SE=5) were true positives, explaining the improved recall rates (Supplementary Table 5). Similarly, there were statistically significant improvements in InDel recall rates (Supplementary Figures 15C and 16C). As expected, adding reads from 5X WGS improved the recall rates the most. The average paired difference in SNV recall rates in WEGS_8P,5X_ was 0.0044 (SE=0.0003) compared to 0.0031 (SE=0.0002) in WEGS_8P,2X_ (Supplementary Table 5).

The change in variant precision rates after adding reads from low-depth WGS differed for SNVs and InDels. We observed slight drops in SNV precision rates in all combinations of multiplexing levels in WES and read depths in WGS. However, the declines were not systematic, i.e. they were present only in part of the samples, in contrast to increases in SNV recall rates which were, on average, much higher and present in all samples (Supplementary Figures 15B and 16B). For example, the lowest average paired difference in SNV precision rates among all WES and WGS combinations was -0.0003 (SE=0.0001) in WEGS_4P,2X_ (Supplementary Table 5). It was the only combination where this paired difference in SNV precision rates reached statistical significance (*P*-value=0.026) (Supplementary Figure 15B). Thus, adding reads from low-depth WGS increased the number of called SNVs by a few dozen, but at the same time, some of these additionally called SNVs were FP, which slightly changed the SNV precision rate in either direction.

Differently from SNVs, all combinations of multiplexing levels in WES and read depths in WGS showed statistically significant improvements in InDel precision rates (P-value < 0.05). In WEGS_8P,2X_, the average paired difference in InDel precision rates was 0.0055 (SE = 0.0009) (Supplementary Table 5), and only 1 out of 16 pairs had a negative paired difference between InDel precision rates after and before adding WGS reads (Supplementary Figure 15). In contrast to SNVs, additional reads from 2X WGS raised the average number of called InDels by 10 (SE=2) and, at the same time, decreased the average number of FPs by 6 (SE=1) in 8-plexing WES.

### 3.6. WEGS enhances WES with millions of variants genome-wide

We compared the variant recall rates in standard no-plexing WES to those in multiplexing WES combined with low-depth WGS (Figures 3A and 3C). The average SNV and InDel recall rates exceeded the corresponding rates in no-plexing WES for most WEGS configurations, except for WEGS_8P,2X_. Both WEGS_4P,2X_ and WEGS_4P,5X_ resulted in higher average SNV recall rates than no-plexing WES: 0.9842 (SE=0.0002, *P*-value=6.4⨉10^-3^) and 0.9852 (SE=0.0001, *P*-value=7.1⨉10^-5^) against 0.9830 (SE=0.0004), respectively (Figure 3A). Among 8-plexing experiments, only WEGS_8P,5X_ resulted in higher average SNV recall rates than no-plexing WES: 0.9847 (SE=0.0001, P-value=5.6⨉10^-4^). Similarly, only WEGS_4P,2X_, WEGS_4P,5X_, and WEGS_8P,5X_ statistically significantly increased average InDel recall rates compared to no-plex WES (Figure 3C). The average InDel recall rate showed a statistically significant increase from 0.9390 (SE=0.0029) in no-plex WES to 0.9493 (SE=0.0029, *P*-value = 0.01), 0.9552 (SE=0.0019, *P*-value = 2.8⨉10^-4^), and 0.9490 (SE=0.0020, *P*-value = 4.2⨉10^-3^) in WEGS_4P,2X_, WEGS_4P,5X_, and WEGS_8P,5X_, respectively. When stratified by the library preparation batch, the average variant recall rates across WEGS remained higher than those in no-plexing WES, except for SNV recall rates in WEGS_8P,2X_ (Supplementary Figures 17E-F, 18E-F). The batch effect in SNV recall rates in WES, described in Section 3.3, also affected WEGS (Supplementary Figure 18D). Despite this, the WEGS_4P,5X_ and WEGS_8P,5X_ had statistically significantly higher SNV recall rates compared to no-plexing WES in both batches, and the increase in WEGS_4P,2X_ was close to statistical significance. (Supplementary Figures 17E-F). There were no statistically significant differences in InDel recall rates between the two batches within the no-plexing WES and each WEGS configuration (Supplementary Figure 18D). But only for WEGS_4P,5X_ the increase in InDel recall rates compared to no-plexing WES was statistically significant in both batches. WEGS_4P,2X_ showed a statistically significant increase only in the first batch. WEGS_8P,2X_ showed a statistically significant increase only in the second batch, and the increase in the first batch was close to a statistical significance (*P*-value=0.092).

**Figure 3.**
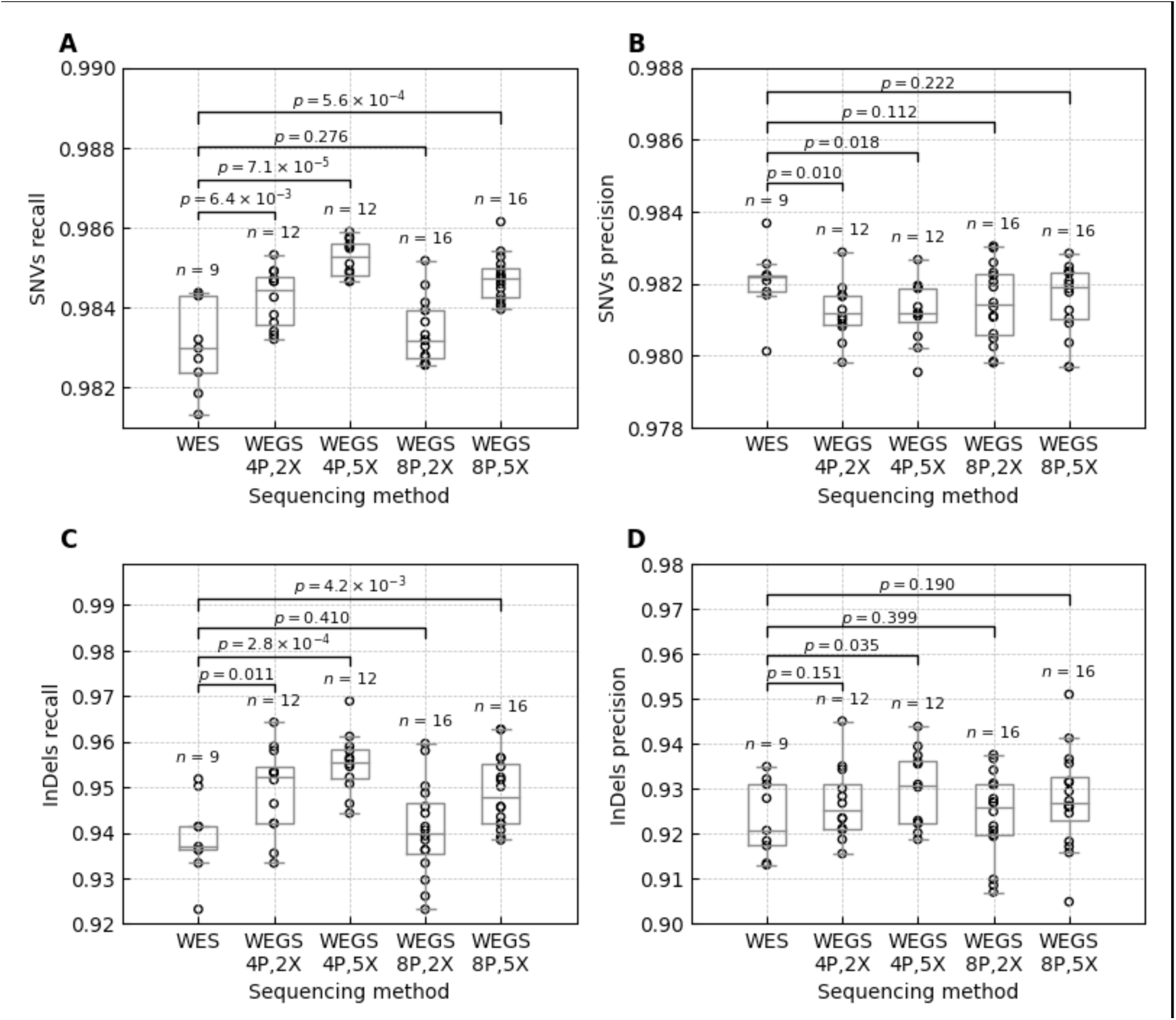
Variant recall and precision rates in no-plexing WES and WEGS. The figure represents variant calls inside the target regions in Agilent V7 capture and the GIAB highconfidence regions. The box bounds the IQR, and Tukey-style whiskers extend to 1.5 × IQR beyond the box. The horizontal line within the box indicates the median value. Open circles are data points corresponding to the individual WES and WEGS. The p-values above each sequencing method pair correspond to the one-tailed Wilcoxon rank-sum test. A) Recall rates of the called SNVs. B) Precision rates of the called SNVs. C) Recall rates of the called InDels D) Precision rates of the called InDels. Supplementary Table 6 shows average values and standard errors.

The variant calling precision rates in no-plexing WES compared to WEGS differed depending on the variant type. The average SNV precision rates in every WEGS configuration were slightly lower than in WES, while average InDel precision rates were higher than in WES (Figures 3B and 3D, Supplementary Table 6). Only drops in average SNV precision rates in WEGS_4P,2X_ and WEGS_4P,5X,_ and an increase in the average InDel precision rate in WEGS_4P,5X_ were statistically significant (Supplementary Table 6). Furthermore, when stratified by the library preparation batch, the decreases in average SNV precision rates in WEGS compared to no-plexing WES were statistically significant only in the second batch (Supplementary Figures 17A-C, Supplementary Table 7). In contrast, WEGS_8P,2X_ and WEGS_8P,5X_ demonstrated an increase in average SNV precision rates compared to no-plexing WES in the first batch. We explain this by the initially lower precision rates in multiplexing WES experiments in the second batch described in Section 3.3. When stratified the average InDel recall rates by the library preparation batches, the average InDel precision rates in WEGS remained higher than in no-plexing WES for all configurations except WEGS_8P,2X_ (Supplementary Figures 18A-C, Supplementary Table 7). However, none of the increases remained statistically significant.

We also compared variant recall and precision rates in WES and WEGS to the 30X WGS, which we generated by downsampling reads from 300X WGS data (see Methods). Average variant recall and precision rates inside regions targeted by WES were higher in 30X WGS compared to WES and WEGS. For SNVs, these differences were below 0.7%, while for InDels, the maximal difference reached 6% (Supplementary Table 8). WEGS_4P,2X_, WEGS_4P,5X_, and WEGS_8P,5X_ were closer to 30X WGS than WES in targeted regions. 30X WGS had no rivals when comparing genome-wide recall and precision rates. On average, it found 1.7-2.5 times more SNVs and 2.5-3.8 times more InDels genome-wide than WEGS (Supplementary Table 9). The average genome-wide SNV and InDel precision rates in WGS were up to 18% and 70% higher than in WEGS, respectively. As expected, WEGS_4P,5X_ and WEGS_8P,5X_ were the closest to the 30X WEGS.

In summary, these results confirm that our WEGS approach eliminates the negative impact of sample multiplexing in WES on variant recall rates in coding regions and brings variant recall rates to the levels of a standard no-plexing WES or higher. Furthermore, these results suggest that WEGS_4P,2X_, WEGS_4P,5X_ and WEGS_8P,5X_ are the closest alternatives to no-plexing WES, as these sequencing strategies demonstrated statistically significant increases in SNV and InDel recall rates and, at the same time, showed increases in InDel precision rates in targeted regions. WEGS has a clear advantage over WES by allowing the assessment of additional ∼2M SNVs and InDels per individual genome-wide.

### 3.7. WEGS correctly assesses variants which genotype imputation misses

Next, we wanted to understand what other benefits low-depth WGS data could bring to multiplexed WES besides removing the negative effects of sample multiplexing. We compared WEGS to array-based genotyping followed by genotype imputation. For each of our three samples, HG002, HG003, and HG004, we emulated the genotyping array data covering 654,013 genetic positions and performed genotype imputation using the TOPMed reference panel consisting of 97,256 diverse genomes. We compared these imputation results to WEGS_4P,2X_ and WEGS_8P,5X_, the closest alternatives to no-plexing WES in targeted regions.

First, we investigated regions targeted by WEGS. SNVs imputed from emulated genotyping array data showed high precision rates (>99%) for all three samples, but imputation missed between 824 to 1,028 SNVs per sample (among them, between 482 to 576 were non-synonymous) compared to WEGS (Supplementary Table 10). For example, in sample HG002, WEGS_8P,5X_ correctly identified 22,390 SNVs on average, and the TOPMed reference panel imputed only 21,458 SNVs which is 938 SNVs less. The difference in the number of correctly identified InDels was even larger: imputation missed around 60% of true InDels (40% recall), while WEGS only missed around 5% (95% recall).

Second, we investigated the number of imputed and sequenced variants genome-wide (Supplementary Table 11). In contrast to the WEGS targeted regions, the genotyping array-based imputation approach outperformed WEGS by the number of correctly identified SNVs: imputation missed 4-5% (95-96% recall), WEGS_4P,2X_ missed 54-65% (35-46% recall), and WEGS_8P,5X_ missed 36-50% (50-64% recall) of true SNVs. The differences in correctly identified InDels were much lower: imputation missed around 61% (39% recall), WEGS_4P,2X_ missed 69%-78% (22-31% recall), and WEGS_8P,5X_ missed 53-67% (33-47% recall) of true InDels.

Third, we looked at how many variants missed or wrongly imputed outside non-protein coding regions can be recovered by WEGS. We grouped TOPMed-imputed variants outside WEGS-targeted non-protein-coding regions into three categories: (1) the number of imputed alleles matches the number of true alleles (i.e. imputation is correct); (2) the number of imputed alleles is less than the number of true alleles; (3) the number of imputed alleles is higher than the number of true alleles. For each of these groups, we looked at the median fold change in alternate AF between the ASJ population and TOPMed. The median fold-change in AF was higher (i.e. AF in ASJ was higher than in TOPMed) when imputation was systematically missing alleles (group 2) and lower when imputation was wrongly imputing extra allele(s) (group 3) (Supplementary Table 12). This result is in line with previous studies^38, 39^, which showed that the imputation accuracy depends on the genetic similarity between the study individual and the reference panel. WEGS_4P,2X_ correctly identified true alleles in 38-46% of variants in group 2 and 89-92% of variants in group 3, while WEGS_8P,5X_ correctly identified true alleles in 55-67% of variants in group 2 and 91-94% in group 3.

Finally, to improve the variant recall in non-coding regions in WEGS, we evaluated the applicability of the GLIMPSE method^31^, developed to impute missing variants from low-depth WGS data. After applying GLIMPSE to WEGS_4P,2X_ and WEGS_8P,5X_ with local reference haplotypes from the 1000 Genomes Project and Human Genome Diversity Project (see Methods), genome-wide SNV recall rates and precision increased drastically. In imputed WEGS_4P,2X_, the average genome-wide SNV recall rate and precision increased from ∼35-46% to ∼81-82% and from ∼80-82% to 87-88%, respectively (Supplementary Tables 11 and 13). In imputed WEGS_8P,5X_, the average genome-wide SNV recall rate and precision increased from ∼50-65% to ∼86-88% and from ∼87-90% to 90-92%, respectively. The genome-wide recall rate and precision also increased for InDels. The SNV recall rate and precision in sequence-based imputation were still smaller than in genotyping array-based imputation. One of the possible explanations is that the state-of-the-art TOPMed reference panel contains >20 times more haplotypes than our local reference panel. To confirm this, we run the genotyping array-based imputation using our local reference panel and the Minimac4 tool^40^. The Minimac4-imputed recall and precision rates for SNVs were similar to the GLIMPSE-imputed WEGS_8P,5X_ (Supplementary Table 14) and slightly higher than the GLIMPSE-imputed WEGS_8P,2X_. The InDel recall rates were higher in the sequence-based imputation compared to genotyping array-based imputation using TOPMed, but lower compared to genotyping array-based imputation using the local reference panel.

In summary, these results showed that WEGS outperforms genotyping array and imputation approach in terms of the number of identified variants, especially InDels, inside protein-coding regions. Outside protein-coding regions, WEGS allows one to discover genetic variants missed by genotyping array-based imputation due to their population specificity. Sequencing-based imputation methods can be applied to WEGS to recover variants missed due to lower depth of coverage outside protein-coding regions. WEGS_8P,5X_ has a clear advantage over WEGS_4P,2X_ outside the protein-coding region due to the higher depth of coverage in the WGS experiment.

### 3.8. WEGS is substantially cheaper than high-depth WES and WGS

We compared costs for WEGS scenarios relative to genotyping arrays, low-depth WGS, 30X WGS and no-plexing 100X WES. Per sample cost estimates for the genotyping array included DNA QC and genotyping using Affymetrix Axiom UKBB array. Sequencing costs per sample were based on current pricing and a scenario of 1,000 samples sequenced on the Illumina NovaSeq 6000, S4 platform. We note that sequencing costs can vary depending on multiple factors, including reagents pricing, flow cell volume and sequencing platform, while genotyping array prices are less affected by sample size.

Our estimates show that the combinations of WEGS_4P,2X_ and WEGS_8P,5X_ are half the price compared to standard 100X WES (no-plexing) and ∼47% of the price of 30X WGS (Table 1). The combination of 5X WGS with 4-plexing WES is slightly more expensive but still 56% of the cost of 30X WGS and 60% of the cost of no-plexing 100X WES. As such, the WEGS scenarios representing the most economical strategies relative to WGS and WES are again the combinations of 2X WGS with 4-plexing WES and 5X WGS with 8-plexing WES. Yet, as shown above, while WEGS_4P,2X_ and WEGS_8P,5X_ show comparable precision and recall in targeted coding regions relative to standard WES, the latter combination is more effective at capturing non-coding variation. As such, we conclude that the most cost-effective WEGS strategy to capture both coding and non-coding variants is 5X WGS with 8-plexing high-depth WES.

**Table 1.** Relative genotyping and sequencing costs per sample given current pricing. Per sample cost estimates for the genotyping array include DNA QC and genotyping using Affymetrix Axiom UKBB array. Sequencing scenarios are based on 1,000 samples sequenced on Illumina NovaSeq 6000, S4 platform. These cost estimates include sample preparation steps from DNA QC (QC, gDNA, high throughput) to Illumina library preparation and capture for Agilent SureSelect XT HS2 V7, sequencing library QC, and Illumina sequencing.

|  | Cost relative to<br>array | Cost relative to<br>2X WGS | Cost relative to<br>5X WGS | Cost relative to<br>30X WGS | Cost relative to<br>100X WES |
| --- | --- | --- | --- | --- | --- |
| Axiom array | 1.00 | 0.52 | 0.33 | 0.08 | 0.09 |
| 2X WGS | 1.91 | 1.00 | 0.63 | 0.15 | 0.17 |
| 5X WGS | 3.04 | 1.59 | 1.00 | 0.24 | 0.26 |
| 30X WGS | 12.48 | 6.52 | 4.10 | 1.00 | 1.08 |
| 100X WES | 11.50 | 6.01 | 3.78 | 0.92 | 1.00 |
| 100X WES 4plex | 5.05 | 2.64 | 1.66 | 0.40 | 0.44 |
| 100X WES 8plex | 3.95 | 2.07 | 1.30 | 0.32 | 0.34 |
| 100X WES 4plex + 2X WGS | 5.80 | 3.03 | 1.91 | **0.47** | **0.50** |
| 100X WES 4plex + 5X WGS | 6.93 | 3.63 | 2.28 | **0.56** | **0.60** |
| 100X WES 8plex + 2X WGS | 4.71 | 2.46 | 1.55 | 0.38 | 0.41 |
| 100X WES 8plex + 5X WGS | 5.84 | 3.05 | 1.92 | **0.47** | **0.51** |

### 3.9. WEGS applied to the study of peripheral artery disease

We applied WEGS to 862 patients diagnosed with PAD and identified 44,747,114 genetic variants (33,505,105 SNVs and 11,242,009 InDels) (Table 2). A total of 12,893,703 of these variants were novel (not described in dbSNP v109.3), from which 63.8% were singletons (carried by one individual). Inside the coding regions, we observed 35.4% synonymous (11,053 per individual), 59.0% non-synonymous (11,636 per individual), 1.1% stop/essential splice (490 per individual), 2.1% frameshift (298 per individual), and 2.3% (371 per individual) inframe genetic variants.

**Table 2.** The number of variants discovered in WEGS sequencing data from 862 patients with peripheral artery disease. This table reports the total number of sequenced variants in the overall patient group and the average number of sequenced variants per individual across different functional classifications. Novel variants were defined as variants not present in dbSNP (version 109.3).

|  | All Individuals (N=862) |  |  | Per Individual |
| --- | --- | --- | --- | --- |
|  | Total | Singletons | Rare (MAF<1%) | Average |
| Depth at non-targeted regions (X) | -- | -- | -- | 4.5 (±0.88) |
| Depth at targeted regions (X) | -- | -- | -- | 114.8 (±4.47) |
| **Total variants** | 44,747,114 | 14,542,812 | 32,497,956 | 2,964,080 |
| SNVs | 33,505,105 | 11,291,220 | 24,576,783 | 2,587,752 |
| InDels | 11,242,009 | 3,226,457 | 7,921,173 | 449,829 |
| **Novel variants** | 12,893,703 | 8,226,183 | 12,712,777 | 26,984 |
| SNVs | 9,363,157 | 5,739,615 | 9,233,592 | 18,922 |
| InDels | 3,530,546 | 2,481,974 | 3,529,310 | 8,062 |
| **Coding variants** | 348,410 | 181,870 | 284,434 | 23,854 |
| Synonymous | 123,337 | 57,968 | 94,275 | 11,053 |
| Non-synonymous | 205,570 | 113,475 | 173,546 | 11,636 |
| Stop/essential splice | 3,967 | 2,325 | 3,426 | 490 |
| Frameshift | 7,375 | 4,484 | 6,520 | 298 |
| Inframe | 8,069 | 3,566 | 6,589 | 371 |
| **Novel coding variants** | 31,821 | 28,448 | 31,713 | 66 |
| Synonymous | 7,680 | 6,920 | 7,675 | 11 |
| Non-synonymous | 19,761 | 17,884 | 19,731 | 31 |
| Stop/essential splice | 1,071 | 865 | 1,045 | 12 |
| Frameshift | 2,166 | 1,897 | 2,159 | 8 |
| Inframe | 1,106 | 872 | 1,067 | 7 |

We evaluated the WEGS ability to capture known loci associated with PAD identified by large-scale GWAS^41^. All lead variants mapping to these loci were present in the PAD WEGS data (Supplementary Table 15). The majority of the lead variants are intergenic, with an average read depth of 13.7. Only 6 out of the 19 lead variants are directly typed onto the Global Screening Array (GSA) 24.v3; demonstrating the WEGS potentials to assess disease-causing variants beyond the genotyping arrays. In addition, we observed that WEGS captured, on average, 4,056 (SE=295) genetic variants within the known PAD loci that are not present in the TOPMed imputation reference panel and, thus, could not be imputed. (Supplementary Table 16). Although the majority of these loci are intergenic, WEGS was able to identify additional missense variants within these regions.

## 4 Discussion

In this work, we propose and evaluate a new sequencing method which we call WEGS, designed to be more economical than WES and WGS. We considered WEGS based on WES (100X) with sample multiplexing, i.e. pooling and sequencing up to 8 samples simultaneously, combined with the low-depth WGS (2-5X). First, we evaluated the effect of sample multiplexing in WES. We demonstrated that an increased number of PCR/optical read duplicates in multiplexing WES experiments leads to the loss of depth of coverage and, consecutively, to a higher number of missed true variants. Second, we showed that although the UMI-aware read deduplication helps improve variant calling recall or precision rates, the improvements are minimal and don’t compensate for the losses due to multiplexing. Third, we demonstrated that combining reads from low-depth WGS and reads from multiplexing WES brings variant calling recall and precision rates in protein-coding regions to the levels of no-plexing WES or above. Specifically, based on our experiments, we recommend using combinations of 2X WGS with 4-plexing WES and 5X WGS with 8-plexing WES as an alternative to standard WES.

When choosing between different WEGS configurations, it is essential to also consider performance outside the protein-coding regions. Specifically, we demonstrated that WEGS allows for the identification of population-specific non-coding genetic variants, which large genotype imputation panels impute less accurately due to differences in allele frequencies between the study population and reference. If there is no available imputation reference panel closely matching the study population, then the 8-plex WES with 5X WGS would be the best option compared to the 4-plex WES with 2X WGS. Also, our cost estimates suggest that WEGS relying on 8-plexing WES and 5X WGS is the most cost-effective configuration and is 2X cheaper than standard no-plex WES and 2.1X cheaper than high-depth WGS. We used this WEGS configuration on 862 samples with PAD to demonstrate the scalability and applicability of the method in a practical setting, assessing almost 3M variations (24,000 in coding regions) per individual genome on average.

The WEGS data processing pipelines build on existing open-source software tools and, thus, does not require time and financial investments in tool development. This work demonstrated how the industry-standard GATK toolset^42^ could be utilized for SNVs and InDels calling and filtering from WEGS data (see Data and Code Availability). Novel genotype imputation methods, such as GLIMPSE^31^, are available for sequencing data and can be applied to WEGS to increase the number of identified non-coding variants further.

Our study has several limitations. First, benchmarking analyses relied on high-confidence variant calls from a GIAB trio. As benchmarking call sets will become available for regions difficult for variant detection (i.e. outside high-confidence regions), it will be interesting to investigate WEGS performance in these regions. Second, our analysis focused on SNVs and InDels only, as WES and low-depth WGS are known to have limited utility for structural variant calling. Third, while our precision and recall estimates were broadly consistent across replicates, we acknowledge that they are based on only 3 individual genomes from a single ancestry. Extension of this work could include an investigation of WEGS performance in individuals from other ancestries. Yet, based on our results and recent work assessing the advantages of low-depth WGS^11^, we expect WEGS to be of particular interest for populations currently underrepresented in public reference panels, enabling the discovery of novel population-specific variants.

We anticipate that WEGS will become a method of choice for studies of the molecular genetic basis of diseases and disease-related traits. Such genetic association studies require many sequenced individuals to reach sufficient statistical power and capacity to detect rare variants. Today, it remains costly to use high-depth WGS; for example, high-depth WGS for 1,000 samples currently costs close to 1 million US dollars, and standard WES can be up to 90% of this figure. As such, a 50% cost reduction when using WEGS will enable high-depth sequencing of up to twice the number of exomes while providing additional information genome-wide. Our cost estimates are based on current pricing, but these relative costs should hold as long as WES reagents costs remain low compared to WGS costs. As such, WEGS should remain competitive until WGS costs become substantially lower than currently. The real impact on association studies will be shown in future studies using WEGS or similar technologies.

## 5 Data and code availability

The generated WES, WGS, and WEGS sequencing data for GIAB samples are available for download at https://datahub-778-pbbb.p.genap.ca/. The variant calling and analysis pipelines and source code are publicly available through version control repositories listed in the table below.

| Name | URL | Description |
| --- | --- | --- |
| WEGS_paper | https://github.com/CERC-Genomic-Medicine/WEGS_paper | Source code for the tables and figures. |
| WEGS_pipelines | https://github.com/CERC-Genomic-Medicine/WEGS_pipelines | Scalable Nextflow pipelines for variant calling in WGS, WES and WEGS using GATK. |
| glimpse_pipeline | https://github.com/CERC-Genomic-Medicine/glimpse_pipeline | A scalable Nextflow pipeline for genotype imputation from low-depth sequencing data using GATK and GLIMPSE. |
| shapeit4_pipeline | https://github.com/CERC-Genomic-Medicine/shapeit_pipeline | A scalable Nextflow pipeline for statistical genotype phasing using SHAPEIT4. |

## 6 Acknowledgments

The authors would like to thank Claire Le Moigne from the McGill Canada Excellence Research Chair Program in Genomic Medicine for administrative support. We also thank the team in Bern who collected the samples and provided administrative support: Györgyi Hamvas, Claudine Andres, Rebecca Scheidegger, Marina Rumenovic, Nathalie Minder. The authors thank the Faculty of Medicine and Biology at Lausanne University, Switzerland, and the Swiss Foundation for Vascular Medicine (Schweizerische Stiftung für Gefässmedizin) for funding. This work was supported by the McGill Canada Excellence Research Chair Program in Genomic Medicine (VM), CFI awards 41012 & 40104 (JR), CFI 33408, CFI MSI 35444 (JR and GB) and Genome Canada (JR). VM holds a Canada Excellence Research Chair. GB is supported by a Canada Research Chair Tier 1 award and a FRQS Distinguished Research Scholar award. The Canadian Center for Computational Genomics (C3G) is supported by the Canadian Government through Genome Canada. We thank Calcul Quebec and the Digital Research Alliance of Canada for computing resources.

## SUPPLEMENTARY FIGURES AND TABLES

**Supplementary Figure 1.**
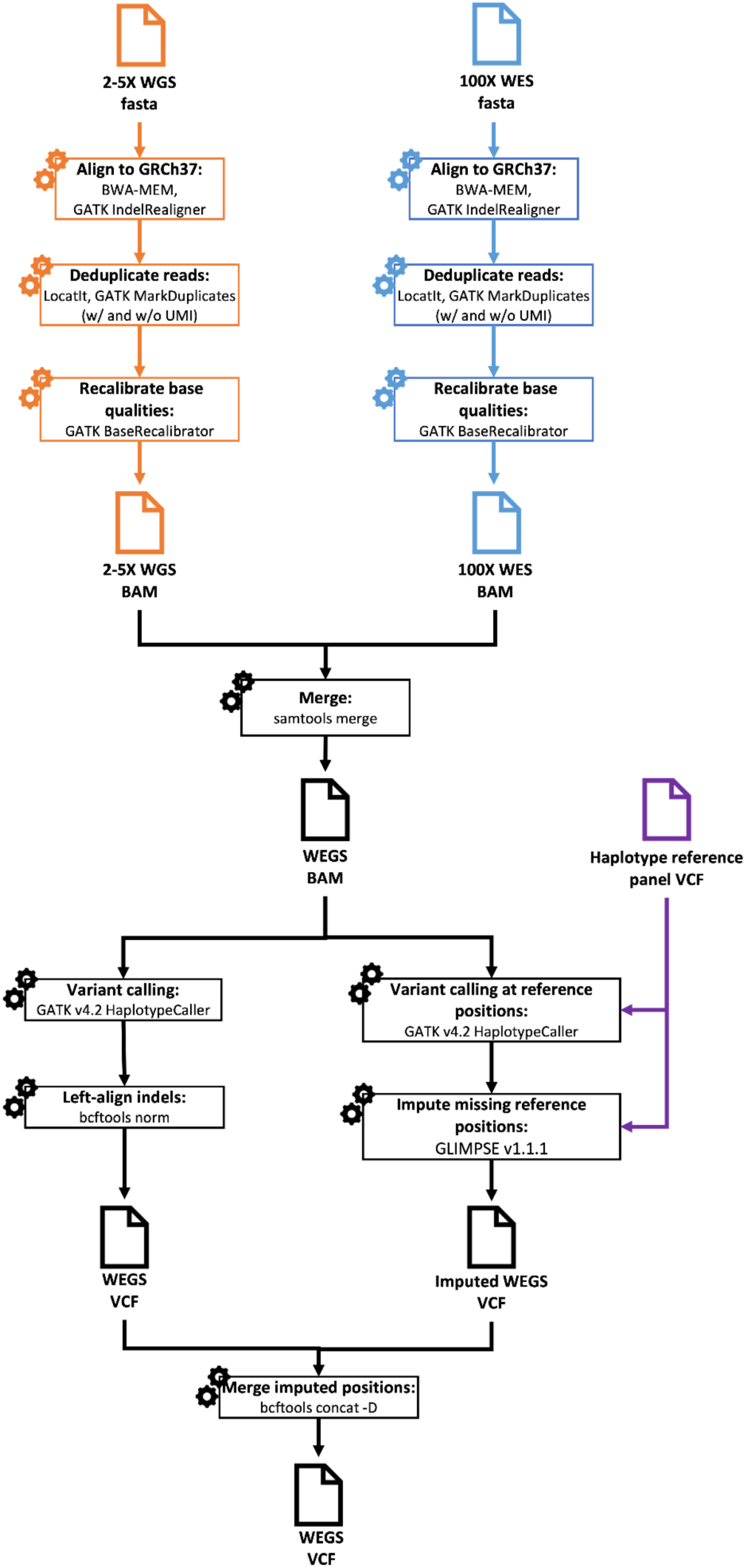
Data processing diagram. WGS and WES alignment, deduplication and base recalibration steps were performed using GATK best practices. WGS and WES BAM files were then merged using samtools. Merged WEGS data was used for variant calling and imputation with GLIMPSE and local haplotype reference panel. The resulting imputed WEGS VCF was then used to merge imputed positions with WEGS data, hence obtaining the final WEGS VCF.

**Supplementary Figure 2.**
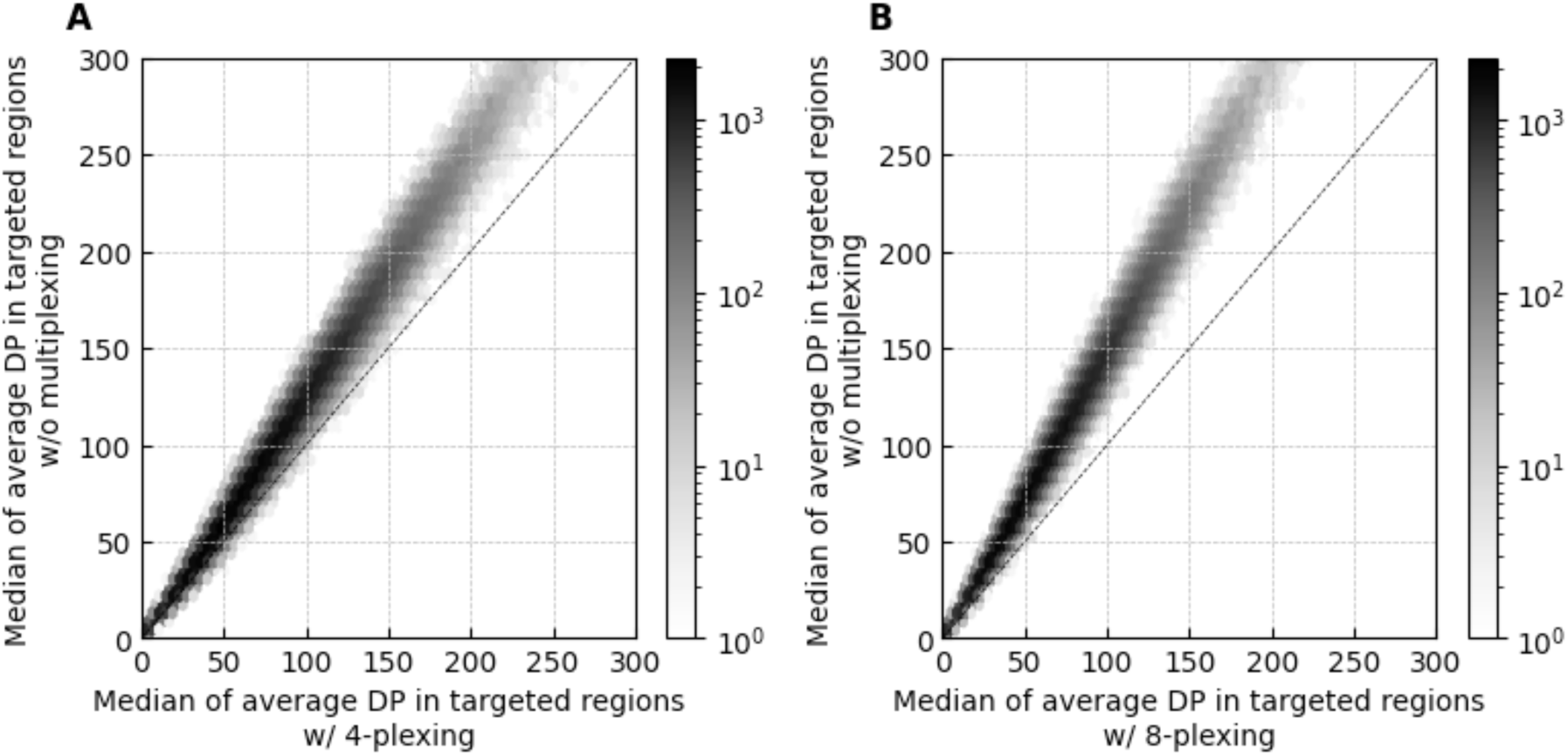
Comparison of average depths of coverage (DP) in individual target regions in experiments without and with multiplexing. We computed the average depth of coverage (DP) for each target region in Agilent V7 capture in each sequenced individual. DP includes only paired mapped reads and base pairs with minimal Phred-scaled mapping and base qualities of 20. Each point corresponds to a single target region. In total, there were 208,817 non-overlapping autosomal regions. The X-axis shows the median of average DPs in the target region across all individuals sequenced in 4-plex (panel A) and 8-plex (panel B) experiments. The Y-axis shows the median of average DPs in the target region across all individuals sequenced without multiplexing. The darker color represents the higher density of the points. The vertical bar on the right of each panel shows the number of points corresponding to each color. The dotted black line corresponds to the 1:1 ratio between DP in the experiments.

**Supplementary Figure 3.**
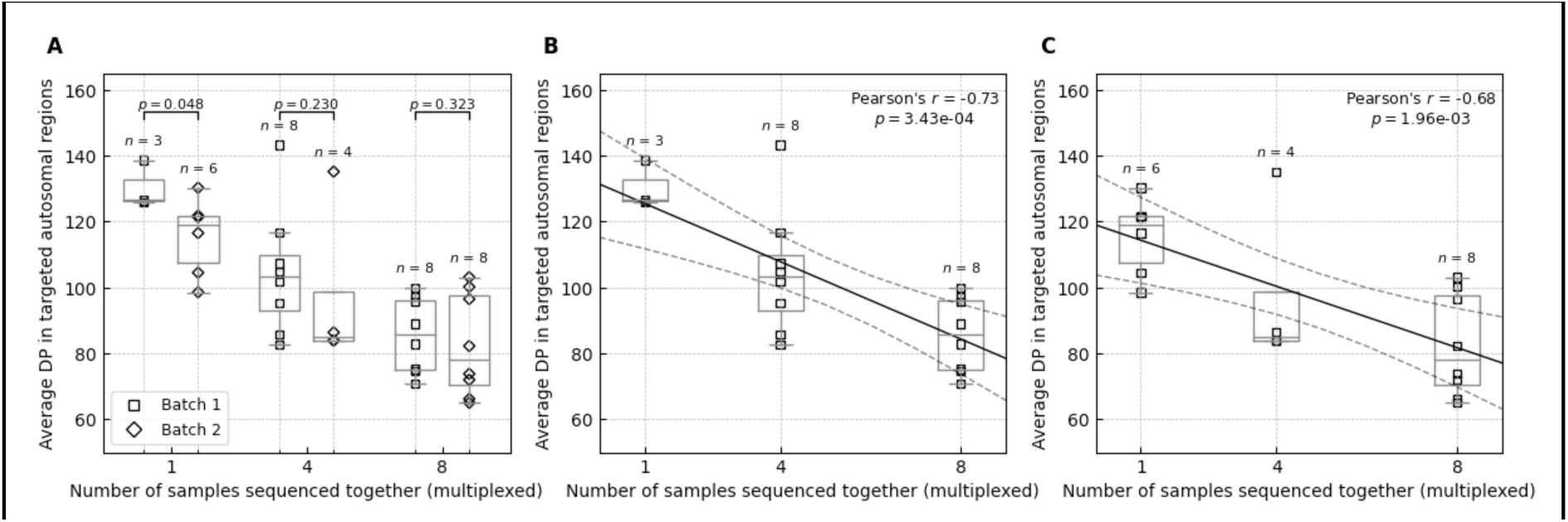
Supplementary Figure 3. Average depths of coverage across all targeted regions in autosomal chromosomes stratified by library preparation batch. The average depth of coverage (DP) was computed across target regions in Agilent V7 capture using paired mapped reads and counting only base pairs with minimal Phred-scaled mapping and base qualities of 20. The solid black line corresponds to the linear regression line, and the dashed black lines correspond to the 95% confidence interval. The box bounds the IQR, and Tukey style whiskers extend to 1.5 ? IQR beyond the box. The horizontal line within the box indicates the median value. Open rectangles and diamonds are data points corresponding to the average DP across individual exome in batches 1 and 2, respectively. A) The DP is stratified by the library preparation batch in experiments without multiplexing, with 4-plexing and 8-plexing experiments. The p-values above each experiment pair correspond to theone-tailed Wilcoxon rank-sum test. B) The DP in the first library preparation batch. C) The DP in the second library preparation batch.

**Supplementary Figure 4.**
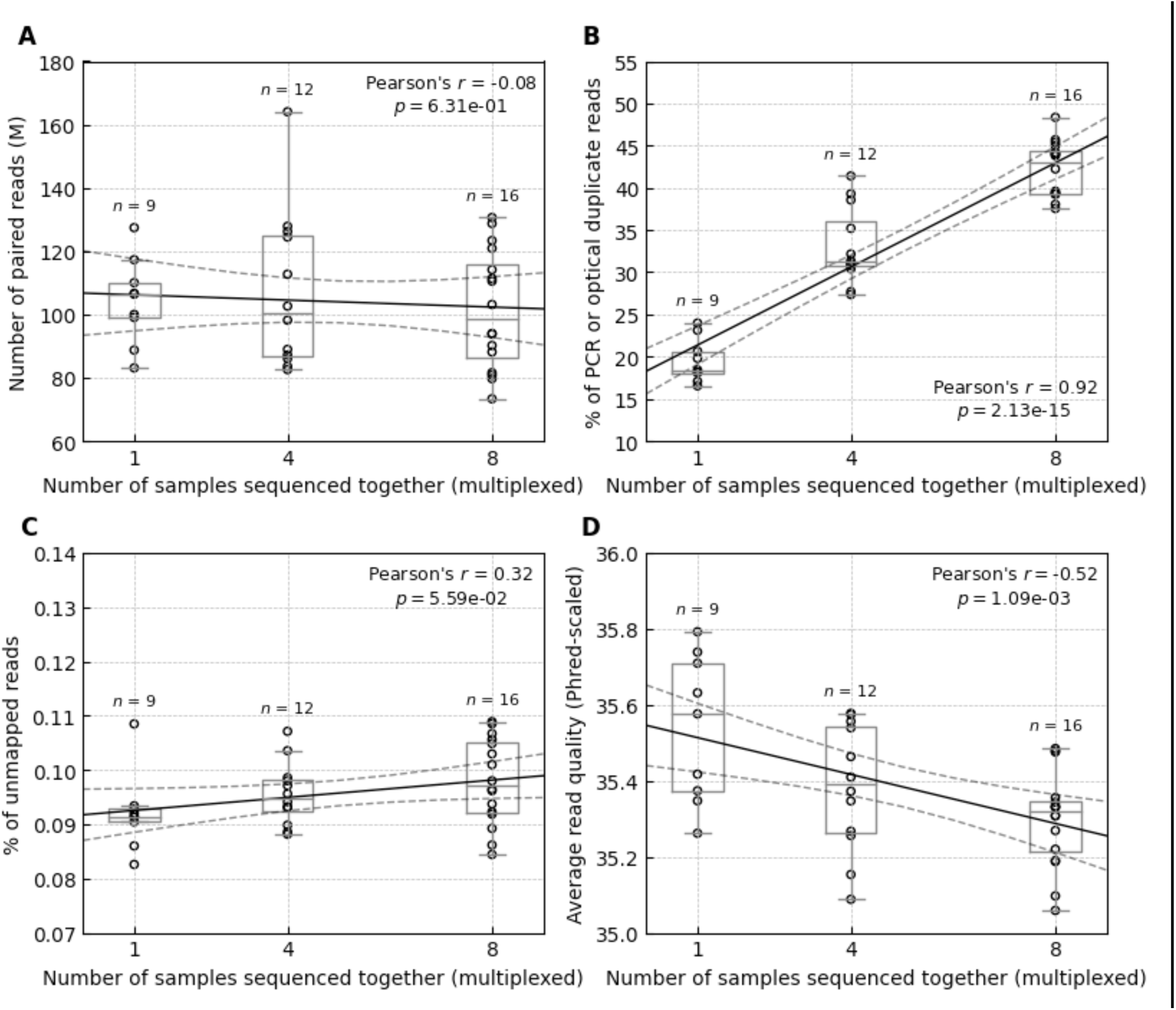
Number of reads in autosomal chromosomes in sequencing experiments with and without multiplexing. The figure shows only paired reads in autosomal chromosomes, excluding reads that are non-primary or supplementary alignments or failed platform/vendor quality checks. The solid black line corresponds to the linear regression line, and the dashed black lines correspond to the 95% confidence interval. The box bounds the IQR, and Tukey-style whiskers extend to 1.5 ⨉ IQR beyond the box. The horizontal line within the box indicates the median value. Open circles are data points corresponding to the sequenced individual exomes. A) Number of paired reads in millions in sequencing experiments without sample multiplexing and when simultaneously sequencing four (4-plex) and eight (8-plex) samples. B) Percent of reads flagged as PCR or optical duplicates. C) Percent of unmapped reads. D) Average Phred-scaled base quality score across all reads in a sequenced sample.

**Supplementary Figure 5.**
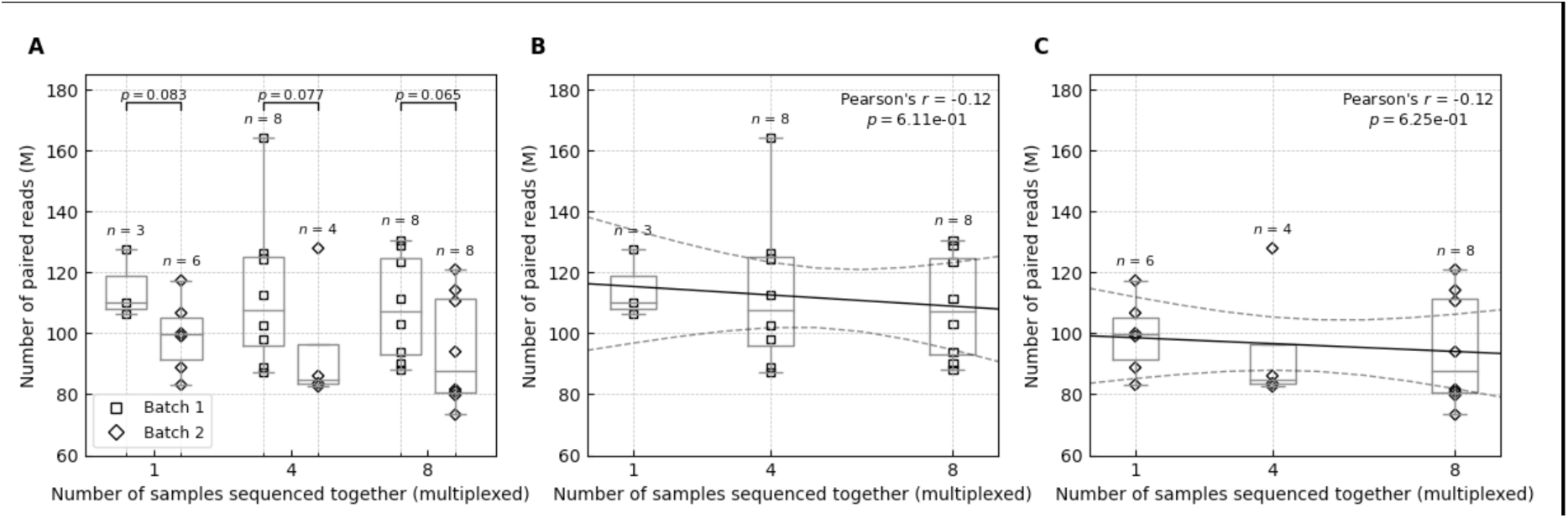
The number of paired reads in autosomal chromosomes stratified by library preparation batch. The figure shows only paired reads in autosomal chromosomes, excluding reads that are non-primary or supplementary alignments or failed platform/vendor quality checks. The solid black line corresponds to the linear regression line, and the dashed black lines correspond to the 95% confidence interval. The box bounds the IQR, and Tukey-style whiskers extend to 1.5 ? IQR beyond the box. The horizontal line within the box indicates the median value. Open rectangles and diamonds are data points corresponding to the number of paired reads across individual exome in batches 1 and 2, respectively. A) The number of paired reads is stratified by the library preparation batch in experiments without multiplexing, with 4-plexing and 8-plexing experiments. The p-values above each experiment pair correspond to the one-tailed Wilcoxon rank-sum test. B) The number of paired reads in the first library preparation batch. C) The number of paired reads in the second library preparation batch.

**Supplementary Figure 6.**
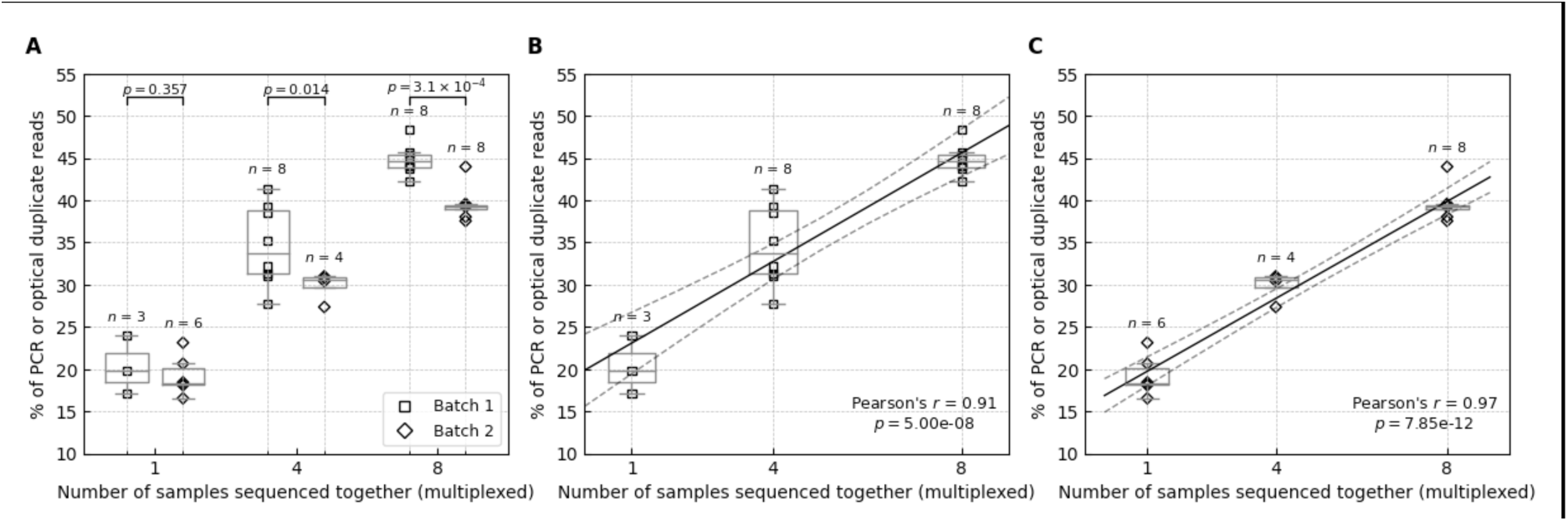
Percent of reads flagged as PCR or optical duplicates in autosomal chromosomes stratified by library preparation batch. The figure shows only paired reads in autosomal chromosomes, excluding reads that are non-primary or supplementary alignments or failed platform/vendor quality checks. The solid black line corresponds to the linear regression line, and the dashed black lines correspond to the 95% confidence interval. The box bounds the IQR, and Tukey-style whiskers extend to 1.5 ? IQR beyond the box. The horizontal line within the box indicates the median value. Open rectangles and diamonds are data points corresponding to the percent of duplicated reads across individual exomes in batches 1 and 2, respectively. A) The percent of duplicated reads is stratified by the library preparation batch in experiments without multiplexing, with 4-plexing and 8-plexing experiments. The p-values above each experiment pair correspond to the one-tailed Wilcoxon rank-sum test. B) The percent of duplicate reads in the first library preparation batch. C) The percent of duplicate reads in the second library preparation batch.

**Supplementary Figure 7.**
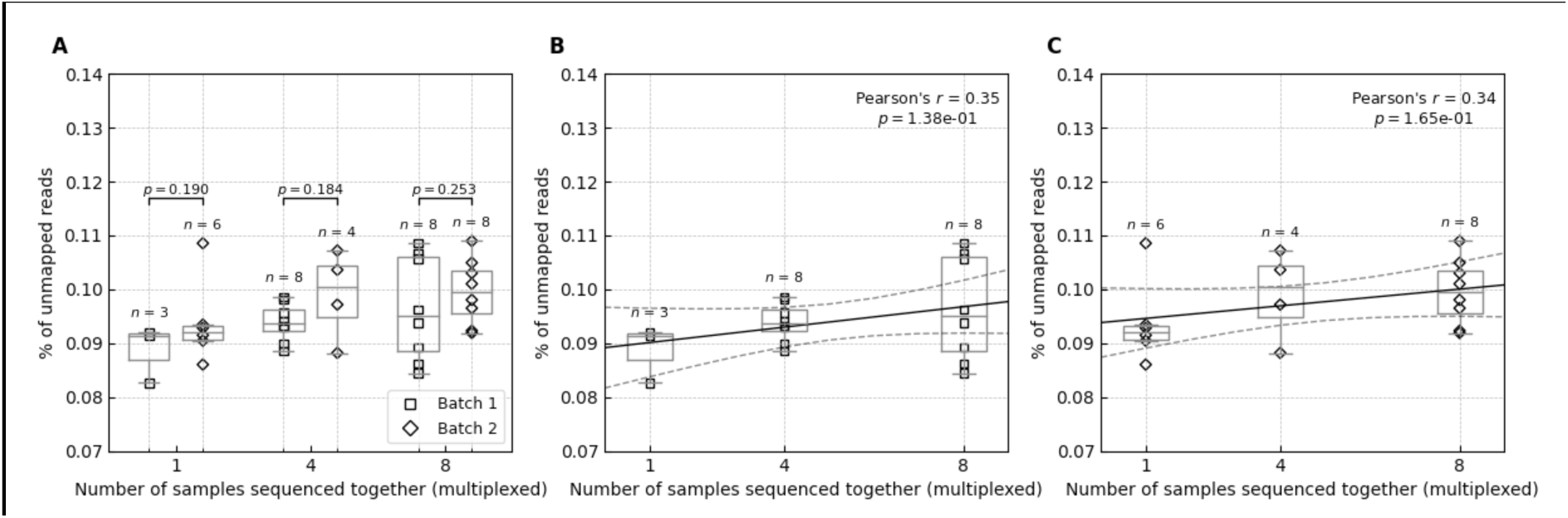
Percent of unmapped reads in autosomal chromosomes stratified by library preparation batch. The figure shows only paired reads in autosomal chromosomes, excluding reads that are non-primary or supplementary alignments or failed platform/vendor quality checks. The solid black line corresponds to the linear regression line, and the dashed black lines correspond to the 95% confidence interval. The box bounds the IQR, and Tukey-style whiskers extend to 1.5 ? IQR beyond the box. The horizontal line within the box indicates the median value. Open rectangles and diamonds are data points corresponding to the percent of unmapped reads across individual exome in batches 1 and 2, respectively. A) The percent of unmapped reads is stratified by the library preparation batch in experiments without multiplexing, with 4-plexing and 8-plexing experiments. The p values above each experiment pair correspond to the one-tailed Wilcoxon rank-sum test. B) The percent of unmapped reads in the first library preparation batch. C) The percent of unmapped reads in the second library preparation batch.

**Supplementary Figure 8.**
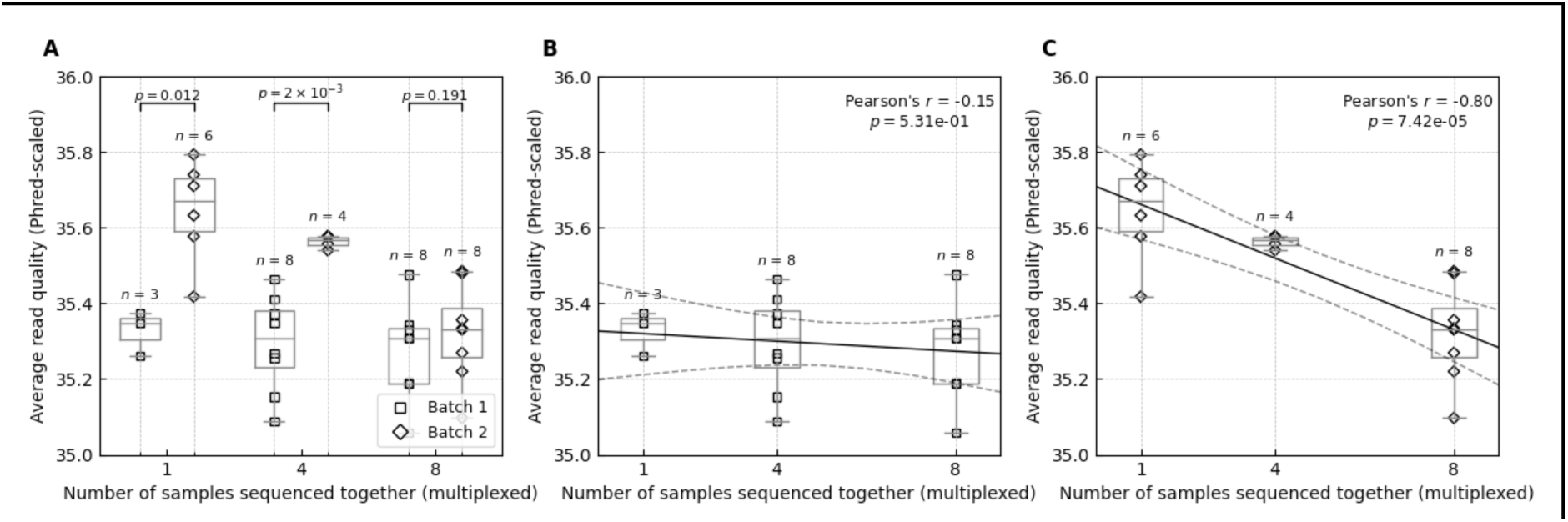
The average quality of reads in autosomal chromosomes stratified by library preparation batch. The figure shows only paired reads in autosomal chromosomes, excluding reads that are non-primary or supplementary alignments or failed platform/vendor quality checks. The average read quality was computed as the average of Phred-scaled base qualities. The solid black line corresponds to the linear regression line, and the dashed black lines correspond to the 95% confidence interval. The box bounds the IQR, and Tukey-style whiskers extend to 1.5 ⨉ IQR beyond the box. The horizontal line within the box indicates the median value. Open rectangles and diamonds are data points corresponding to the average read quality across individual exome in batches 1 and 2, respectively. A) The average read quality is stratified by the library preparation batch in experiments without multiplexing, with 4-plexing and 8-plexing experiments. The p-values above each experiment pair correspond to the one-tailed Wilcoxon rank-sum test. B) The average read quality in the first library preparation batch. C) The average read quality in the second library preparation batch.

**Supplementary Figure 9.**
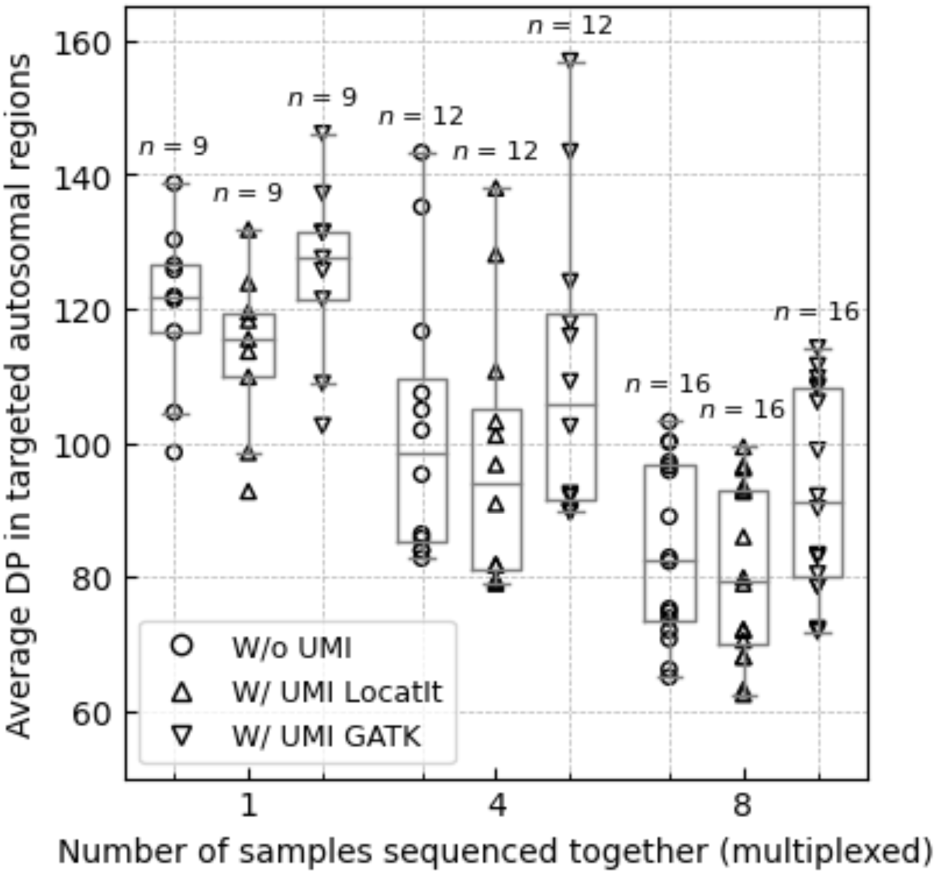
Average depths of coverage across all targeted regions in autosomal chromosomes processed with and without UMI-aware deduplication. The average depth of coverage (DP) was computed across target regions in Agilent V7 capture using paired mapped reads and counting only base-pairs with minimal Phred-scaled mapping and base qualities of 20. The box bounds the IQR and Tukey-style whiskers extend to a maximum of 1.5 × IQR beyond the box. The horizontal line within the box indicates median value. Open circles, up-pointing and down-pointing triangles are data points corresponding to the average DP across individual exome processed without, with LocatIt and GATK’s UmiAwareMarkDuplicatesWithMateCigar UMI-aware deduplication, respectively.

**Supplementary Figure 10.**
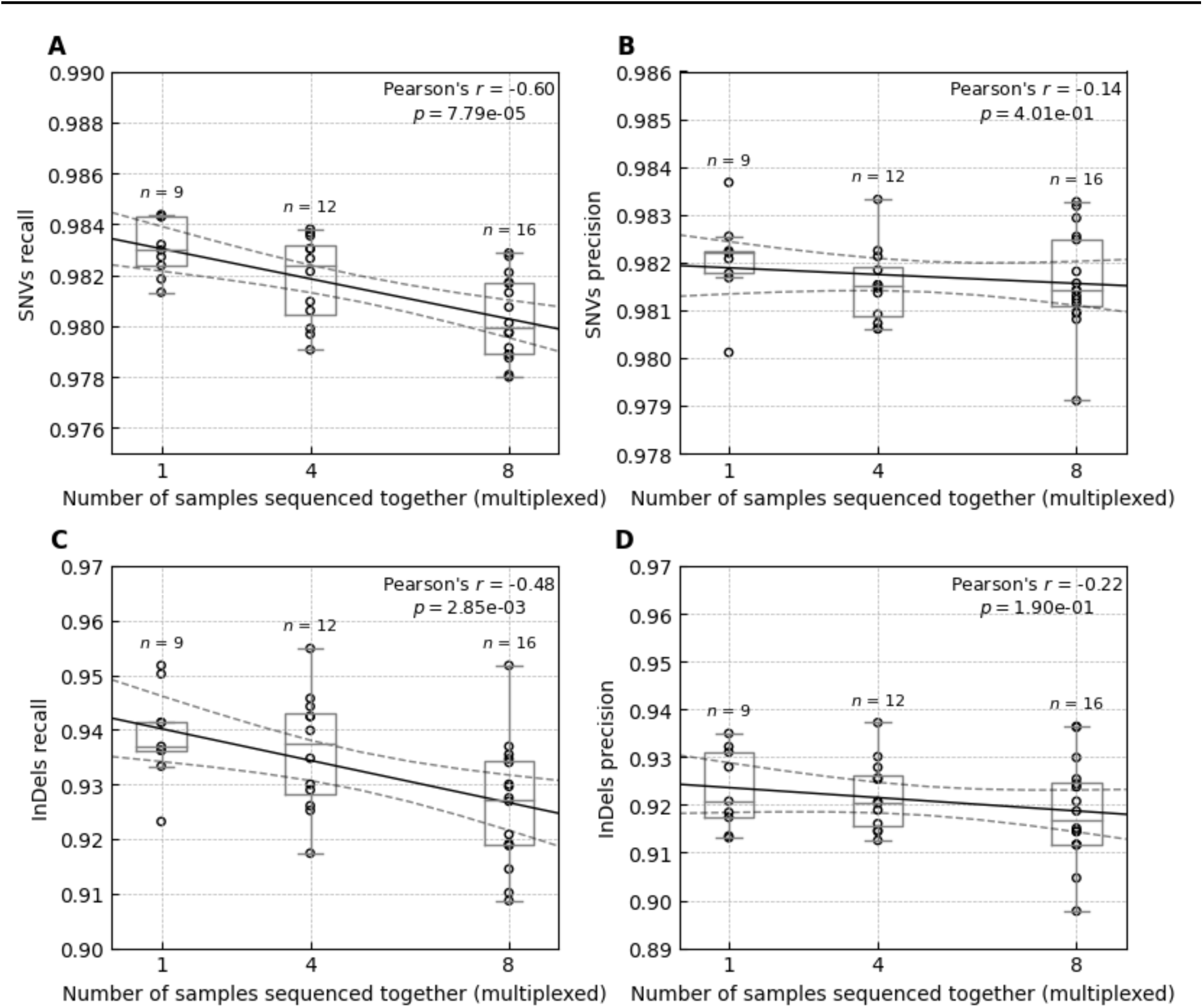
Recall and precision of the SNVs and InDels called in sequencing experiments without and with multiplexing. The figure represents variant calls inside the target regions in Agilent V7 capture and the GIAB high-confidence regions. The solid black line corresponds to the linear regression line, and the dashed black lines correspond to the 95% confidence interval. The box bounds the IQR, and Tukey-style whiskers extend to 1.5 × IQR beyond the box. The horizontal line within the box indicates the median value. Open circles are data points corresponding to the sequenced individual exomes. A) Recall rates of the called SNVs. B) Precision of the called SNVs. C) Recall rates of the called InDels. D) Precision of the called InDels.

**Supplementary Figure 11.**
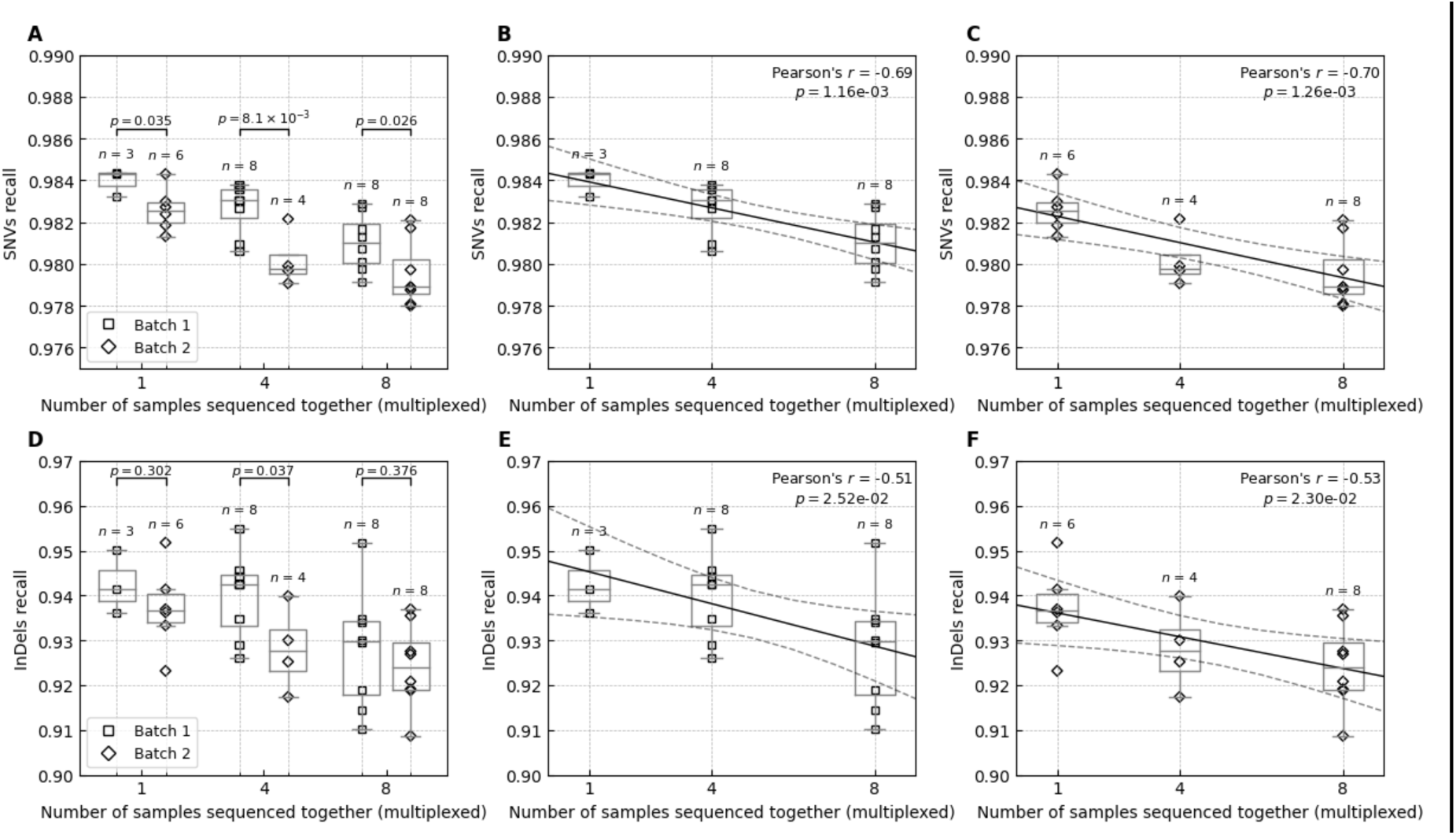
The recall of the SNVs and InDels called in sequencing experiments without and with multiplexing stratified by library preparation batch. The figure represents variant calls inside the target regions in Agilent V7 capture and the GIAB high-confidence regions. The solid black line corresponds to the linear regression line, and the dashed black lines correspond to the 95% confidence interval. The box bounds the IQR, and Tukey-style whiskers extend to 1.5 × IQR beyond the box. The horizontal line within the box indicates the median value. Open rectangles and diamonds are data points corresponding to the recall across individual exome in batches 1 and 2, respectively. A) The recall of SNVs is stratified by the library preparation batch in experiments without multiplexing, with 4-plexing and 8-plexing experiments. The p-values above each experiment pair correspond to the one-tailed Wilcoxon rank-sum test. B) The recall of SNVs in the first library preparation batch. C) The recall of SNVs in the second library preparation batch. D) The recall of InDels is stratified by the library preparation batch in experiments without multiplexing, with 4-plexing and 8-plexing experiments. The p-values above each experiment pair correspond to the one-tailed Wilcoxon rank-sum test. E) The recall of InDels in the first library preparation batch. F) The recall of InDels in the second library preparation batch.

**Supplementary Figure 12.**
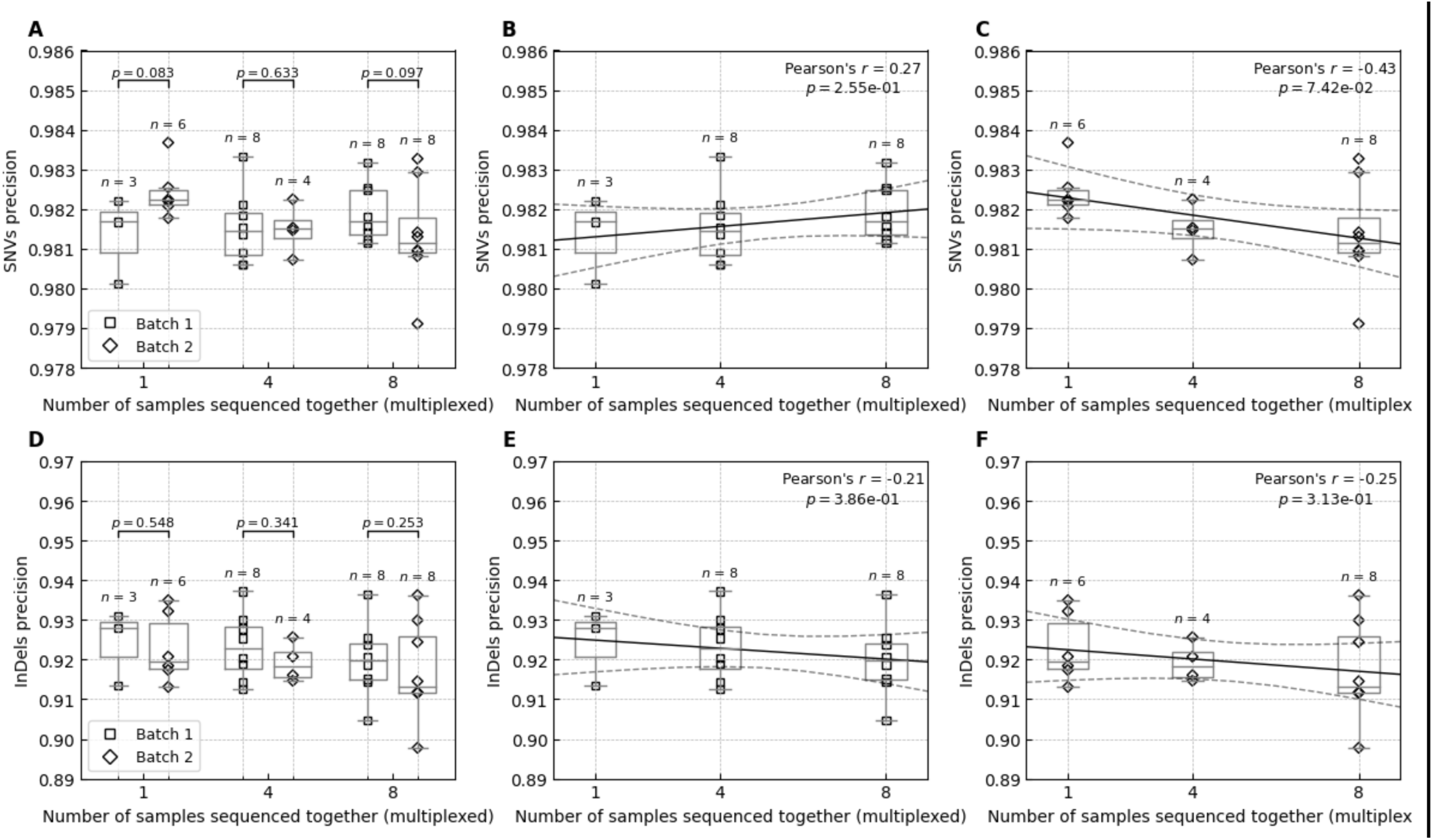
The precision of the SNVs and InDels called in sequencing experiments without and with multiplexing stratified by library preparation batch. The figure represents variant calls inside the target regions in Agilent V7 capture and the GIAB high-confidence regions. The solid black line corresponds to the linear regression line, and the dashed black lines correspond to the 95% confidence interval. The box bounds the IQR, and Tukey-style whiskers extend to 1.5 × IQR beyond the box. The horizontal line within the box indicates the median value. Open rectangles and diamonds are data points corresponding to the precision across individual exome in batches 1 and 2, respectively. A) The precision of SNVs is stratified by the library preparation batch in experiments without multiplexing, with 4-plexing and 8-plexing experiments. The p-values above each experiment pair correspond to the one-tailed Wilcoxon rank-sum test. B) The precision of SNVs in the first library preparation batch. C) The precision of SNVs in the second library preparation batch. D) The precision of InDels is stratified by the library preparation batch in experiments without multiplexing, with 4-plexing and 8-plexing experiments. The p-values above each experiment pair correspond to the one-tailed Wilcoxon rank-sum test. E) The precision of InDels in the first library preparation batch. F) The precision of InDels in the second library preparation batch.

**Supplementary Figure 13.**
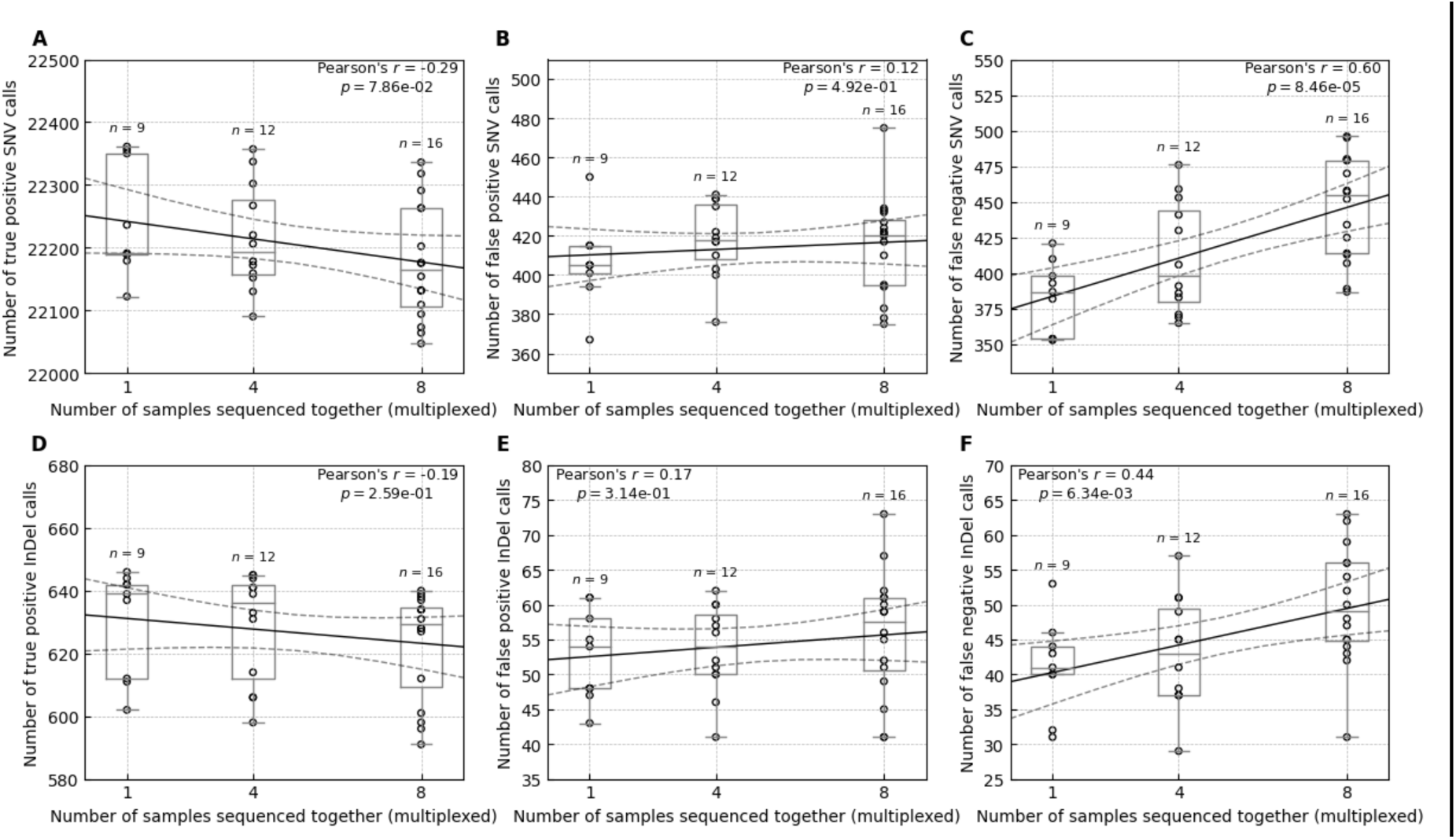
The number of SNV and InDel calls in sequencing experiments without and with multiplexing. The figure represents variant calls inside the target regions in Agilent V7 capture and the GIAB high-confidence regions. The solid black line corresponds to the linear regression line, and the dashed black lines correspond to 95% confidence interval. The box bounds the IQR, and Tukey-style whiskers extend to 1.5 × IQR beyond the box. The horizontal line within the box indicates the median value. A) Number of true positive (TP) SNV calls in sequencing experiments without sample multiplexing and when simultaneously sequencing four (4-plex) and eight (8-plex) samples. B) Number of false positive (FP) SNV calls. C) Number of false negative (FN) SNV calls. D) Number of TP InDel calls. E) Number FP InDel calls. F) Number of FN InDel calls.

**Supplementary Figure 14.**
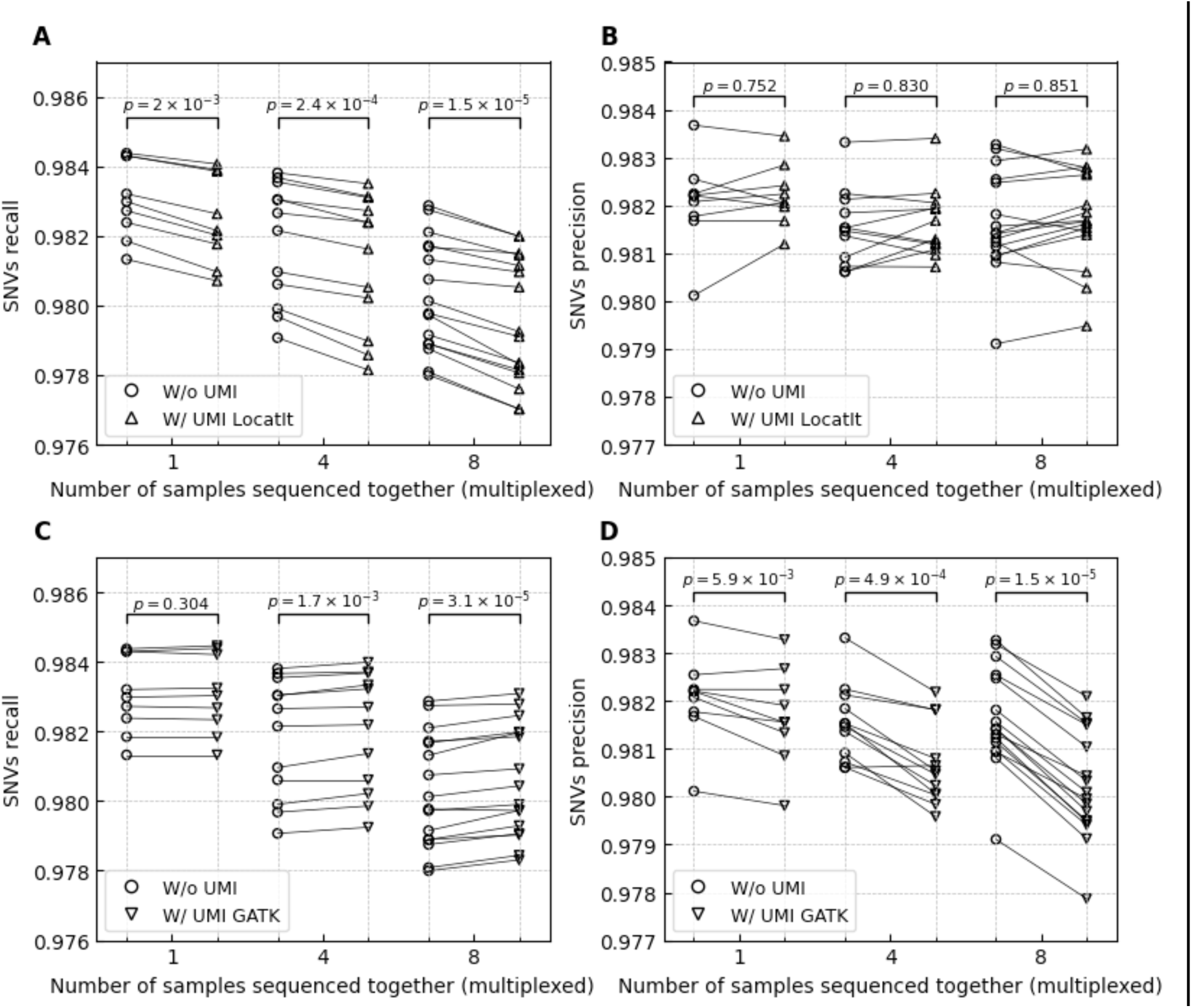
Recall and precision of the single nucleotide variations (SNVs) in sequencing experiments without and with UMI-aware read deduplication. The figure represents SNV calls inside the target regions in Agilent V7 capture and the GIAB high-confidence regions. Open circles, up-pointing and down-pointing triangles are data points corresponding to the recall and precision in individual exomes processed without, with LocatIt and GATK’s UmiAwareMarkDuplicatesWithMateCigar UMI-aware deduplication, respectively. The solid black lines connect pairs of individual exomes with the same underlying sequencing data (i.e. same sequenced sample) but different deduplication approaches. The p-values above experiments with varying levels of multiplexing correspond to the one-tailed Wilcoxon signed-rank test between UMI agnostic and UMI-aware deduplication. A) Recall rates of the called SNVs without UMI-aware compared to UMI-aware deduplication using LocatIt. B) Precision of the called SNVs without UMI-aware compared to UMI-aware deduplication using LocatIt. C) Recall rates of the called SNVs without UMI-aware compared to UMI-aware deduplication using GATK’s UmiAwareMarkDuplicatesWithMateCigar. D) Precision of the called SNVs without UMI-aware compared to UMI-aware deduplication using GATK’s UmiAwareMarkDuplicatesWithMateCigar.

**Supplementary Figure 15.**
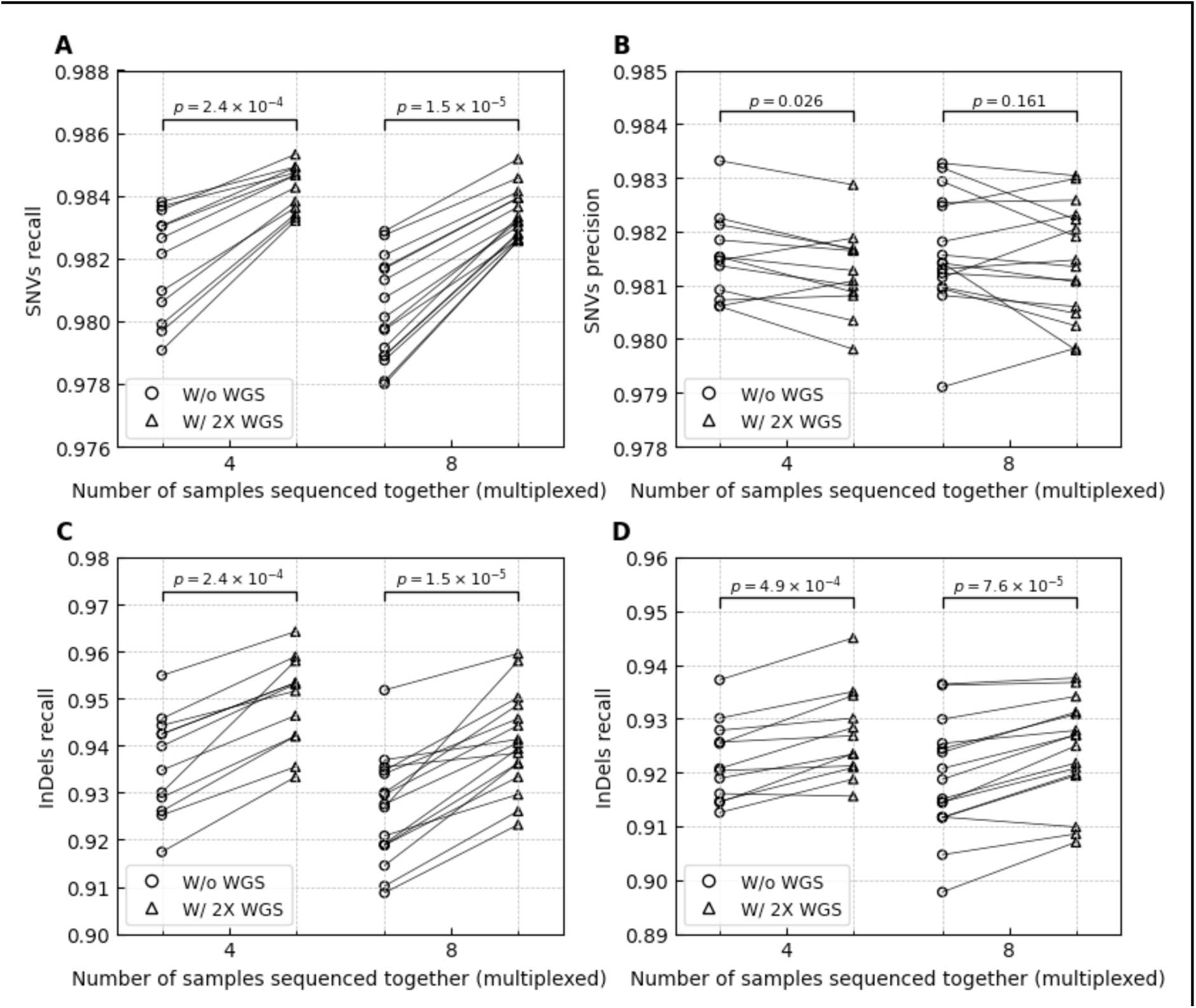
Variant recall and precision rates in WES experiments with multiplexing before and after adding 2X WGS data. The figure represents variant calls inside the target regions in Agilent V7 capture and the GIAB high-confidence regions. Open circles and up-pointing triangles are data points corresponding to the recall and precision in individual multiplexed WES before and after adding 2X WGS data, respectively. The solid black lines connect pairs of individual exomes with the same underlying WES data (i.e. same sequenced sample). The p-values above experiments with varying levels of multiplexing correspond to the one-tailed Wilcoxon signed-rank test between UMI agnostic and UMI-aware deduplication. A) Recall rates of the called SNVs with and without 2X WGS. B) Precision rates of the called SNVs with and without 2X WGS. C) Recall rates of the called InDels with and without 2X WGS. D) Precision rates of the called InDels with and without 2X WGS.

**Supplementary Figure 16.**
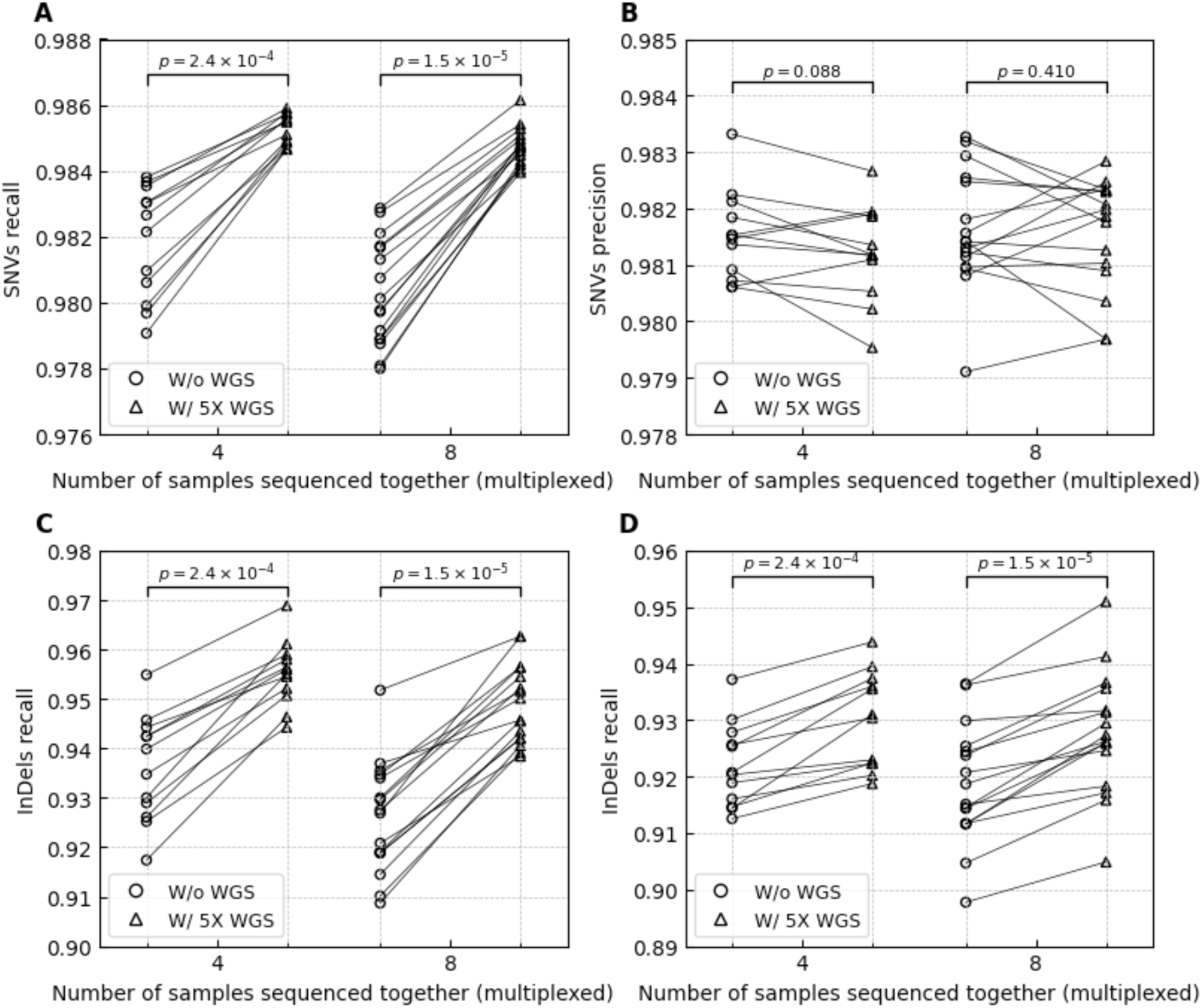
Variant recall and precision rates in WES experiments with multiplexing before and after adding 5X WGS data. The figure represents variant calls inside the target regions in Agilent V7 capture and the GIAB high-confidence regions. Open circles and up-pointing triangles are data points corresponding to the recall and precision in individual multiplexed WES before and after adding 5X WGS data, respectively. The solid black lines connect pairs of individual exomes with the same underlying WES data (i.e. same sequenced sample). The p-values above experiments with varying levels of multiplexing correspond to the one-tailed Wilcoxon signed-rank test between UMI agnostic and UMI-aware deduplication. A) Recall rates of the called SNVs with and without 5X WGS. B) Precision rates of the called SNVs with and without 5X WGS. C) Recall rates of the called InDels with and without 5X WGS. D) Precision rates of the called InDels with and without 5X WGS.

**Supplementary Figure 17.**
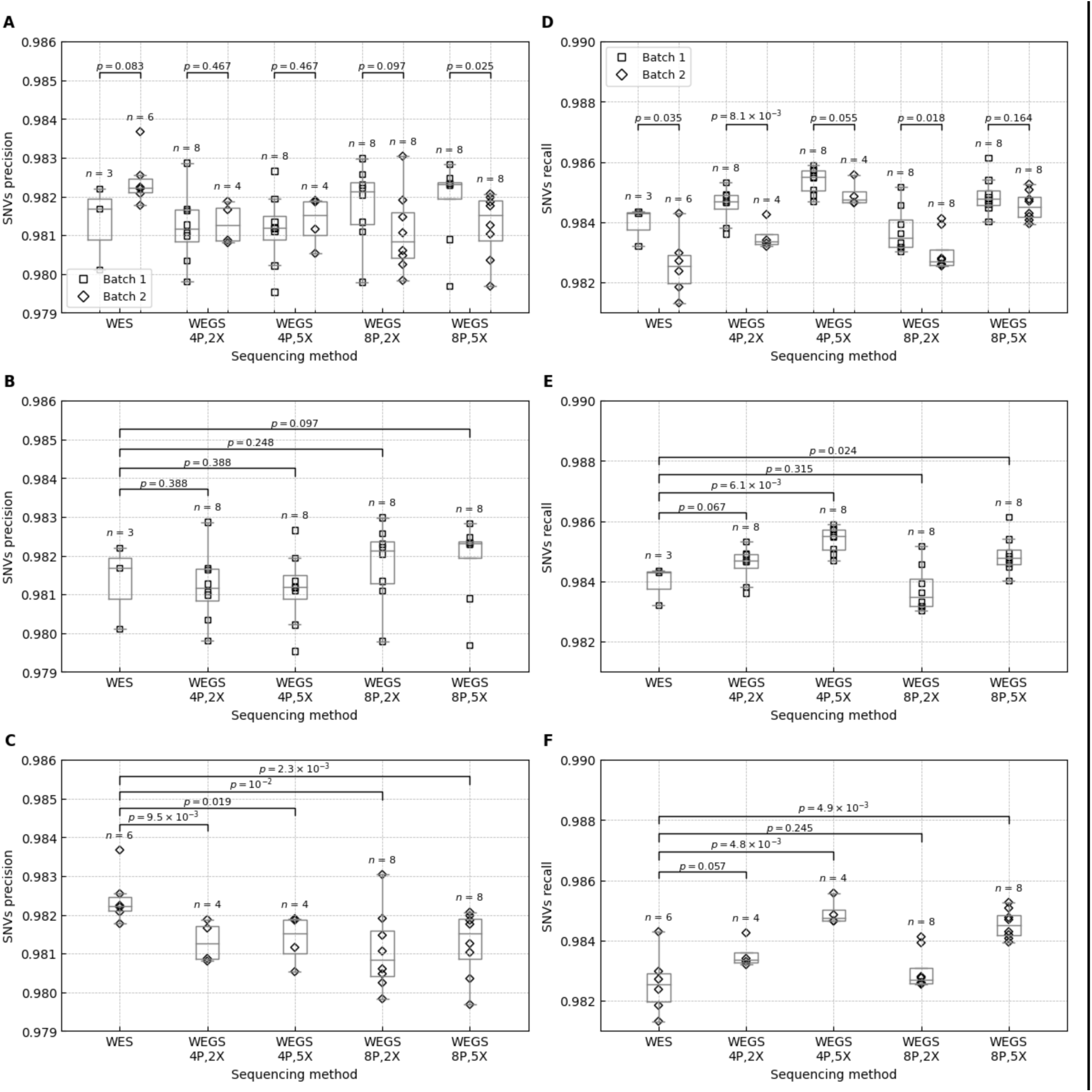
SNVs calling precision and recall rates in no-plexing WES compared to WEGS stratified by library preparation batch. The figure represents SNV calls inside the target regions in Agilent V7 capture and the GIAB high-confidence regions. The box bounds the IQR, and Tukey-style whiskers extend to 1.5 × IQR beyond the box. The horizontal line within the box indicates the median value. Open rectangles and diamonds are data points corresponding to the individual WES and WEGS in batches 1 and 2, respectively. The p-values above each pair of batches or sequencing methods correspond to the one-tailed Wilcoxon rank-sum test. A) Precision rates of the called SNVs in batches 1 and 2. B) Precision rates of the called SNVs in batch 1. C) Precision rates of the called SNVs in batch 2. D) Recall rates of the called SNVs in batches 1 and 2. E) Recall rates of the called SNVs in batch 1. F) Recall rates of the called SNVs in batch 2. Supplementary Table 7 shows average values and standard errors.

**Supplementary Figure 18.**
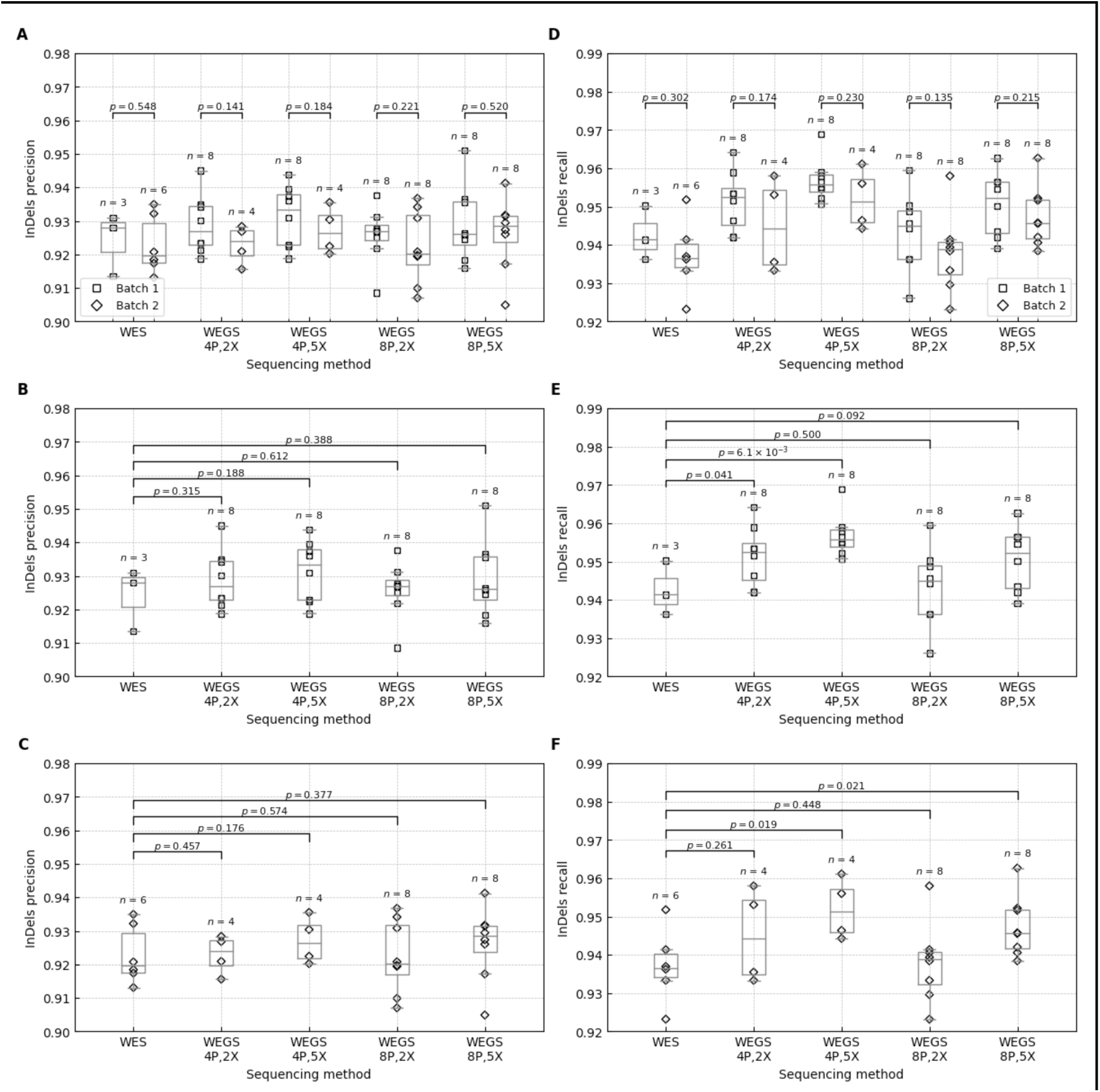
InDel calling precision and recall rates in no-plexing WES compared to WEGS stratified by library preparation batch. The figure represents InDel calls inside the target regions in Agilent V7 capture and the GIAB high-confidence regions. The box bounds the IQR, and Tukey-style whiskers extend to 1.5 × IQR beyond the box. The horizontal line within the box indicates the median value. Open rectangles and diamonds are data points corresponding to the individual WES and WEGS in batches 1 and 2, respectively. The p-values above each pair of batches or sequencing methods correspond to the one-tailed Wilcoxon rank-sum test. A) Precision rates of the called InDels in batches 1 and 2. B) Precision rates of the called InDels in batch 1. C) Precision rates of the called InDels in batch 2. D) Recall rates of the called InDels in batches 1 and 2. E) Recall rates of the called InDels in batch 1. F) Recall rates of the called InDels in batch 2. Supplementary Table 7 shows average values and standard errors.

**Supplementary Table 1.** The average changes in read properties after UMI-aware read deduplication steps relative to the UMI agnostic approach.

| Number of samples<br>sequenced<br>together<br>(multiplexed) | UMI-aware<br>deduplication tool | Average difference compared to UMI agnostic approach |  |  |  |
| --- | --- | --- | --- | --- | --- |
|  |  | % of QC fail reads<br>(SE) | % of PCR/optical<br>duplicates (SE) | % of unmapped<br>reads (SE) | Avg. Phred-scaled<br>read quality (SE) |
| 1 | LocatIt | 6.45 (0.16) | -1.20 (0.04) | -0.068 (0.00223) | 2.60 (0.12) |
| 1 | GATK | 0.00 (0.00) | -0.36 (0.01) | 0.003 (0.00003) | 0.35 (0.06) |
| 4 | LocatIt | 5.19 (0.16) | -1.42 (0.04) | -0.069 (0.00159) | 2.87 (0.05) |
| 4 | GATK | 0.00 (0.00) | -0.39 (0.01) | 0.003 (0.00005) | 0.49 (0.05) |
| 8 | LocatIt | 4.38 (0.10) | -1.56 (0.03) | -0.069 (0.00175) | 2.70 (0.14) |
| 8 | GATK | 0.00 (0.00) | -0.40 (0.01) | 0.003 (0.00004) | 0.57 (0.03) |

**Supplementary Table 2.** Variant calling in whole exome sequencing experiments with and without multiplexing. The table represents variant calls inside the target regions in Agilent V7 capture and the GIAB high-confidence regions. TP - true positives, FP - false positives, FN - false negatives.

| Number of samples<br>sequenced together<br>(multiplexed) | SNVs |  |  |  |  |  | InDels |  |  |  |  |  |
| --- | --- | --- | --- | --- | --- | --- | --- | --- | --- | --- | --- | --- |
|  | N<br>(SE) | TP<br>(SE) | FP<br>(SE) | FN<br>(SE) | Recall<br>(SE) | Precision<br>(SE) | N<br>(SE) | TP<br>(SE) | FP<br>(SE) | FN<br>(SE) | Recall<br>(SE) | Precision<br>(SE) |
| 1 | 22,648<br>(33) | 22,241<br>(30) | 406<br>(7) | 384<br>(8) | 0.9830<br>(0.0004) | 0.9821<br>(0.0003) | 683<br>(7) | 631<br>(6) | 53<br>(2) | 41<br>(2) | 0.9390<br>(0.0029) | 0.9232<br>(0.0028) |
| 4 | 22,632<br>(28) | 22,214<br>(24) | 418<br>(6) | 411<br>(11) | 0.9818<br>(0.0005) | 0.9815<br>(0.0002) | 682<br>(7) | 629<br>(5) | 54<br>(2) | 43<br>(2) | 0.9360<br>(0.0031) | 0.9220<br>(0.0021) |
| 8 | 22,592<br>(28) | 22,177<br>(23) | 415<br>(6) | 446<br>(9) | 0.9803<br>(0.0004) | 0.9816<br>(0.0003) | 679<br>(6) | 623<br>(4) | 56<br>(2) | 50<br>(2) | 0.9261<br>(0.0028) | 0.9186<br>(0.0027) |

**Supplementary Table 3.**
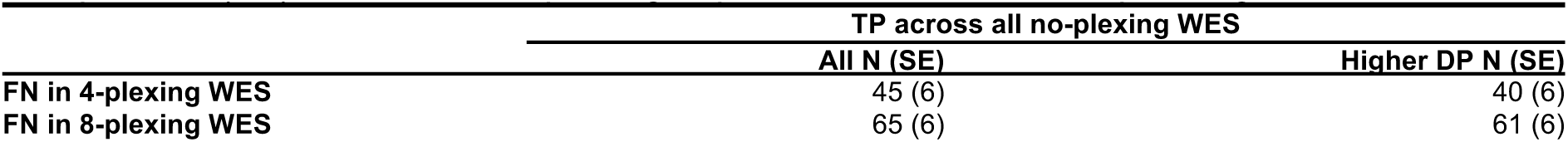
The average number of SNVs missed in multiplexing experiments but correctly identified across all no-plexing experiments. For each multiplexing experiment, we computed the number of false negative (FN) SNV calls that were true positive (TP) in all three no-plexing experiments for the corresponding individual.

|  | TP across all no-plexing WES |  |
| --- | --- | --- |
|  | All N (SE) | Higher DP N (SE) |
| FN in 4-plexing WES | 45 (6) | 40 (6) |
| FN in 8-plexing WES | 65 (6) | 61 (6) |

**Supplementary Table 4.** The average changes in SNV calling in whole exome sequencing experiments with UMI-aware read deduplication relative to the UMI agnostic approach. The table represents SNV calls inside the target regions in Agilent V7 capture and the GIAB high-confidence regions. The star symbols represent statistically significant differences when using a one-tailed Wilcoxon signed-rank test: * - P-value < 0.05, ** - P-value < 0.01, *** - P-value < 0.001.

| Number of samples<br>sequenced together<br>(multiplexed) | UMI-aware<br>deduplication<br>tool | N (SE) | TP (SE) | FP (SE) | FN (SE) | Recall (SE) | Precision (SE) |
| --- | --- | --- | --- | --- | --- | --- | --- |
| 1 | LocatIt | -17 (3) ** | -14 (1) ** | -4 (4) | 14 (1) ** | -0.0006 (0.0001) ** | 0.0002 (0.0002) |
| 1 | GATK | 9 (3) ** | 0 (0) | 8 (2) ** | -0 (0) | 0.0000 (0.0000) | -0.0004 (0.0001)** |
| 4 | LocatIt | -16 (3) *** | -13 (2) *** | -3 (3) | 13 (2) *** | -0.0006 (0.0001)*** | 0.0001 (0.0001) |
| 4 | GATK | 24 (3) *** | 4 (1) ** | 20 (3) *** | -4 (1) ** | 0.0002 (<0.0001) ** | -0.0009 (0.0001)*** |
| 8 | LocatIt | -20 (4) *** | -17 (2) *** | -2 (3) | 17 (2) *** | -0.0008 (0.0001)*** | 0.0001 (0.0001) |
| 8 | GATK | 39 (2) *** | 6 (1) *** | 33 (2) *** | -6 (1) *** | 0.0003 (<0.0001)*** | -0.0014 (0.0001)*** |

**Supplementary Table 5.** The average changes in SNVs and InDels calling in whole exome sequencing experiments when adding additional whole genome sequencing reads relative to pure whole exome sequencing experiments. The table represents SNV and InDel calls inside the target regions in Agilent V7 capture and the GIAB high-confidence regions. The star symbols represent statistically significant differences when using a one-tailed Wilcoxon signed-rank test: * - P-value < 0.05, ** - P- value < 0.01, *** - P-value < 0.001.

| Number of samples sequenced together (multiplexed) | WGS DP | SNVs |  |  |  |  |  | InDels |  |  |  |  |  |
| --- | --- | --- | --- | --- | --- | --- | --- | --- | --- | --- | --- | --- | --- |
|  |  | N (SE) | TP (SE) | FP (SE) | FN (SE) | Recall (SE) | Precision (SE) | N (SE) | TP (SE) | FP (SE) | FN (SE) | Recall (SE) | Precision (SE) |
| 4 | 2X | 62 (8) *** | 54 (7) *** | 8 (3) * | -54 (7) *** | 0.0024 (0.0003) *** | -0.0003 (0.0001) * | 6 (1) ** | 9 (1) *** | -3 (1) ** | -9 (1) *** | 0.0133 (0.0015) *** | 0.0049 (0.0010) *** |
| 8 | 2X | 76 (6) *** | 70 (5) *** | 6 (4) | -70 (5) *** | 0.0031 (0.0002) *** | -0.0002 (0.0002) | 6 (2) *** | 10 (1) *** | -3 (1) *** | -10 (1) *** | 0.0146 (0.0017) *** | 0.0055 (0.0009) *** |
| 4 | 5X | 85 (9) *** | 77 (9) *** | 8 (4) * | -77 (9) *** | 0.0034 (0.0004) *** | -0.0003 (0.0002) | 8 (1) *** | 13 (1) *** | -5 (1) *** | -13 (1) *** | 0.0192 (0.0020) *** | 0.0080 (0.0013) *** |
| 8 | 5X | 104 (8) *** | 100 (7) *** | 3 (5) | -100 (7) *** | 0.0044 (0.0003) *** | -0.0001 (0.0002) | 10 (2) *** | 15 (1) *** | -6 (1) *** | -15 (1) *** | 0.0229 (0.0018) *** | 0.0092 (0.0011) *** |

**Supplementary Table 6.** Average variant recall and precision rates in no-plexing WES and WEGS. The table represents variant calls inside the target regions in Agilent V7 capture and the GIAB high-confidence regions. The star symbols represent statistically significant differences between WES and WEGS when using a one-tailed Wilcoxon rank-sum test: * - P- value < 0.05, ** - P-value < 0.01, *** - P-value < 0.001. WEGS values in bold font are higher than the corresponding values in WES.

| Sequencing method | Number of samples sequenced together (multiplexed) | WGS DP | SNVs |  | InDels |  |
| --- | --- | --- | --- | --- | --- | --- |
|  |  |  | Recall (SE) | Precision (SE) | Recall (SE) | Precision (SE) |
| WES | 1 | — | 0.9830 (0.0004) | 0.9821 (0.0003) | 0.9390 (0.0029) | 0.9232 (0.0028) |
| WEGS<sub>4P,2X</sub> | 4 | 2 | **0.9842 (0.0002)\*\*** | 0.9812 (0.0002)* | **0.9493 (0.0028)\*** | **0.9269 (0.0024)** |
| WEGS<sub>4P,5X</sub> | 4 | 5 | **0.9852 (0.0001)\*\*\*** | 0.9812 (0.0002)* | **0.9552 (0.0019)\*\*\*** | **0.9300 (0.0024)\*** |
| WEGS<sub>8P,2X</sub> | 8 | 2 | **0.9834 (0.0002)** | 0.9814 (0.0003) | **0.9407 (0.0026)** | **0.9240 (0.0024)** |
| WEGS<sub>8P,5X</sub> | 8 | 5 | **0.9847 (0.0001)\*\*\*** | 0.9816 (0.0002) | **0.9490 (0.0020)\*\*** | **0.9277 (0.0027)** |

**Supplementary Table 7.**
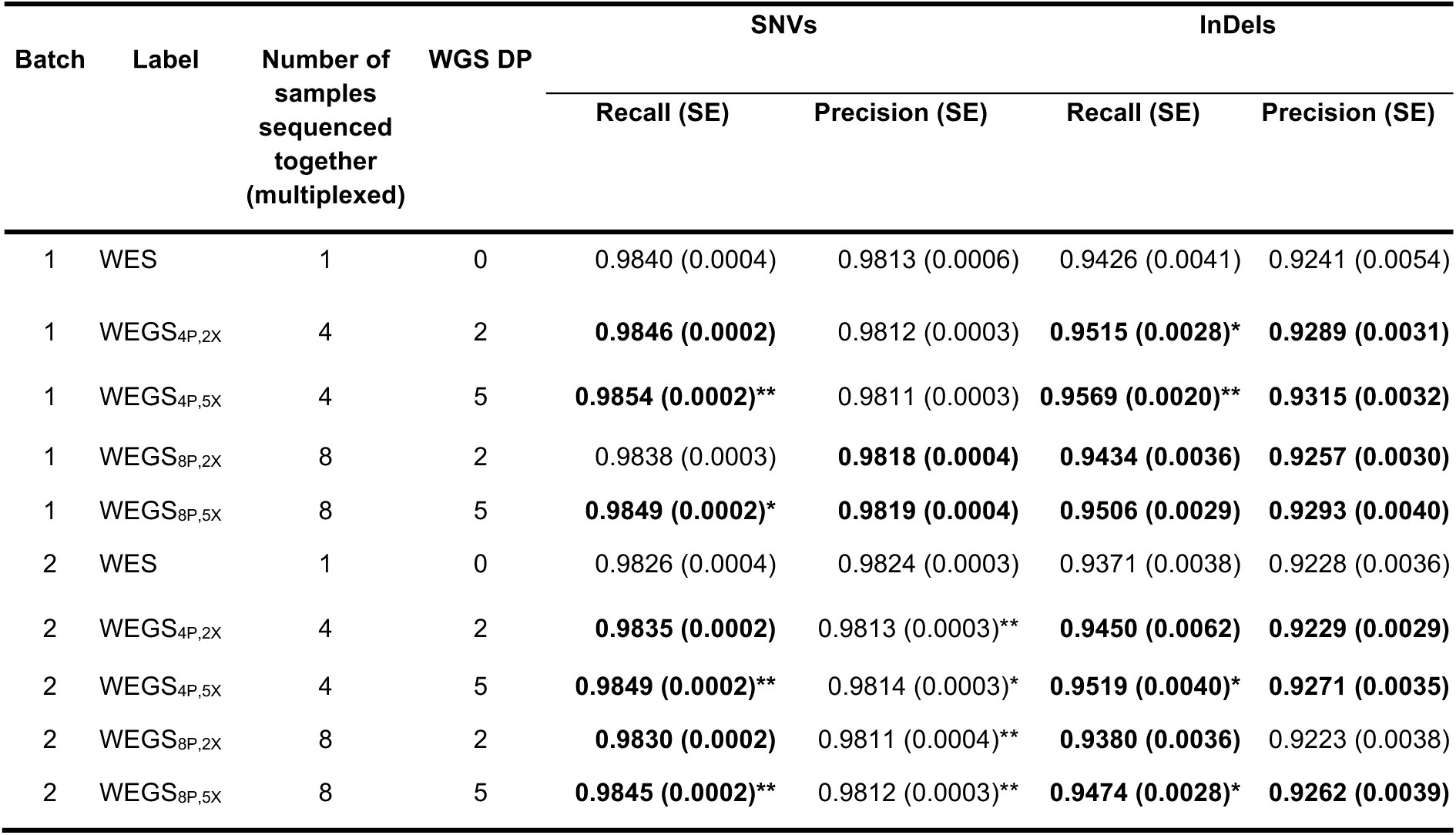
Average variant recall and precision rates in no-plexing WES and WEGS stratified by library preparation batch. The star symbols represent statistically significant differences between WES and WEGS in the same batch when using a one-tailed Wilcoxon rank-sum test: * - P-value < 0.05, ** - P-value < 0.01, *** - P-value < 0.001. WEGS values in bold font are higher than the corresponding values in WES in the same batch.

| Batch | Label | Number of samples sequenced together (multiplexed) | WGS DP | SNVs |  | InDels |  |
| --- | --- | --- | --- | --- | --- | --- | --- |
|  |  |  |  | Recall (SE) | Precision (SE) | Recall (SE) | Precision (SE) |
| 1 | WES | 1 | 0 | 0.9840 (0.0004) | 0.9813 (0.0006) | 0.9426 (0.0041) | 0.9241 (0.0054) |
| 1 | WEGS<sub>4P,2X</sub> | 4 | 2 | **0.9846 (0.0002)** | 0.9812 (0.0003) | **0.9515 (0.0028)\*** | **0.9289 (0.0031)** |
| 1 | WEGS<sub>4P,5X</sub> | 4 | 5 | **0.9854 (0.0002)\*\*** | 0.9811 (0.0003) | **0.9569 (0.0020)\*\*** | **0.9315 (0.0032)** |
| 1 | WEGS<sub>8P,2X</sub> | 8 | 2 | 0.9838 (0.0003) | **0.9818 (0.0004)** | **0.9434 (0.0036)** | **0.9257 (0.0030)** |
| 1 | WEGS<sub>8P,5X</sub> | 8 | 5 | **0.9849 (0.0002)\*** | **0.9819 (0.0004)** | **0.9506 (0.0029)** | **0.9293 (0.0040)** |
| 2 | WES | 1 | 0 | 0.9826 (0.0004) | 0.9824 (0.0003) | 0.9371 (0.0038) | 0.9228 (0.0036) |
| 2 | WEGS<sub>4P,2X</sub> | 4 | 2 | **0.9835 (0.0002)** | 0.9813 (0.0003)** | **0.9450 (0.0062)** | **0.9229 (0.0029)** |
| 2 | WEGS<sub>4P,5X</sub> | 4 | 5 | **0.9849 (0.0002)\*\*** | 0.9814 (0.0003)* | **0.9519 (0.0040)\*** | **0.9271 (0.0035)** |
| 2 | WEGS<sub>8P,2X</sub> | 8 | 2 | **0.9830 (0.0002)** | 0.9811 (0.0004)** | **0.9380 (0.0036)** | 0.9223 (0.0038) |
| 2 | WEGS<sub>8P,5X</sub> | 8 | 5 | **0.9845 (0.0002)\*\*** | 0.9812 (0.0003)** | **0.9474 (0.0028)\*** | **0.9262 (0.0039)** |

**Supplementary Table 8.**
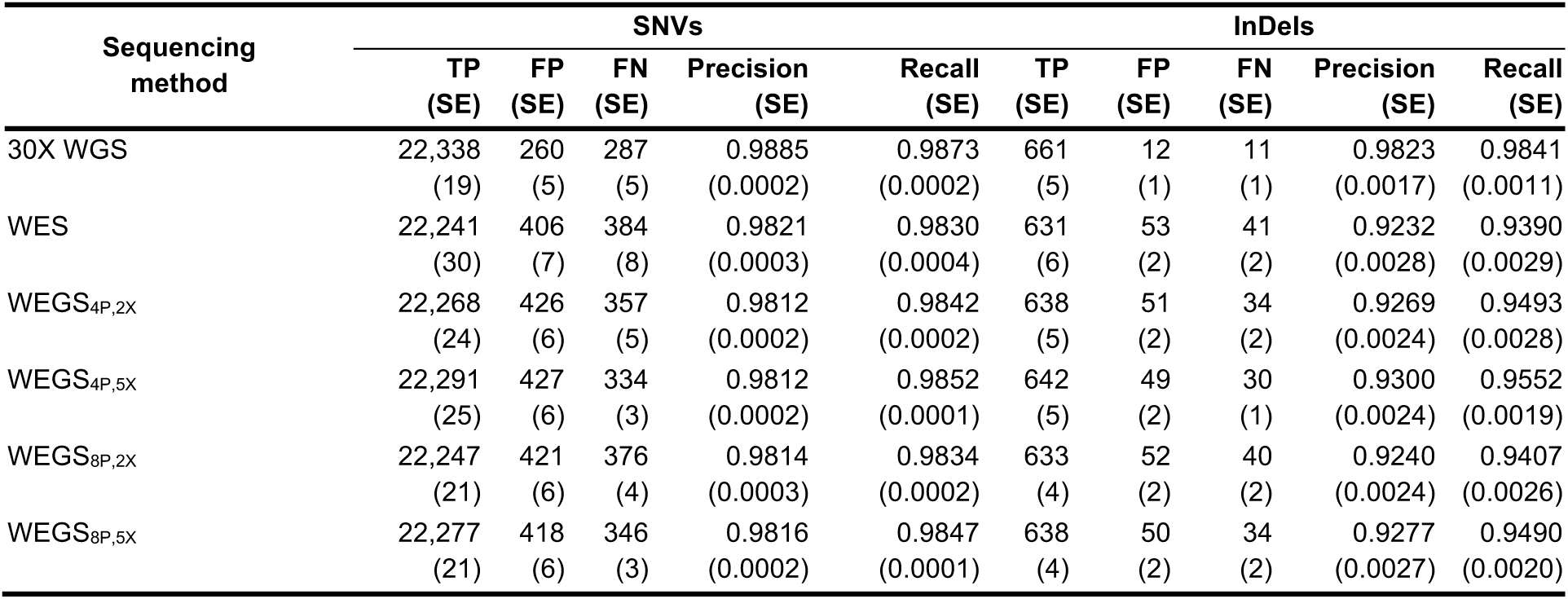
Average variant recall and precision rates in 30X WGS, WES, and WEGS. The table represents variant calls inside the target regions in Agilent V7 capture and the GIAB high-confidence regions.

**Supplementary Table 9.** Average genome-wide variant recall and precision rates in 30X WGS and WEGS. The table represents variant calls in genetic regions overlapping with the GIAB high-confidence regions genome-wide.

| Sequencing method | SNV |  |  |  |  | InDel |  |  |  |  |
| --- | --- | --- | --- | --- | --- | --- | --- | --- | --- | --- |
|  | TP (SE) | FP (SE) | FN (SE) | Precision (SE) | Recall (SE) | TP (SE) | FP (SE) | FN (SE) | Precision (SE) | Recall (SE) |
| 30X WGS | 3,309,667<br>(4,295) | 15,268<br>(174) | 26,097<br>(82) | 0.9954<br>(0.0001) | 0.9922<br>(0.0000) | 500,728<br>(2,382) | 5,605<br>(35) | 9,757<br>(111) | 0.9889<br>(0.0001) | 0.9809<br>(0.0002) |
| WEGS<sub>4P,2X</sub> | 1,333,840<br>(44,603) | 303,452<br>(4,673) | 2,001,925<br>(48,123) | 0.8137<br>(0.0033) | 0.4000<br>(0.0138) | 132,918<br>(5,359) | 56,441<br>(1,663) | 377,567<br>(6,801) | 0.7010<br>(0.0025) | 0.2607<br>(0.0112) |
| WEGS<sub>4P,5X</sub> | 1,909,331<br>(58,921) | 240,392<br>(3,054) | 1,426,434<br>(62,566) | 0.8869<br>(0.0043) | 0.5726<br>(0.0183) | 201,593<br>(7,908) | 68,206<br>(1,221) | 308,892<br>(9,249) | 0.7457<br>(0.0041) | 0.3954<br>(0.0166) |
| WEGS<sub>8P,2X</sub> | 1,326,086<br>(39,124) | 307,039<br>(4,029) | 2,008,327<br>(42,308) | 0.8109<br>(0.0029) | 0.3979<br>(0.0121) | 131,783<br>(4,687) | 56,193<br>(1,460) | 378,097<br>(6,013) | 0.7001<br>(0.0022) | 0.2588<br>(0.0098) |
| WEGS<sub>8P,5X</sub> | 1,914,743<br>(51,710) | 240,678<br>(2,681) | 1,419,670<br>(54,999) | 0.8870<br>(0.0038) | 0.5745<br>(0.0161) | 201,995<br>(6,927) | 68,102<br>(1,067) | 307,885<br>(8,174) | 0.7463<br>(0.0036) | 0.3967<br>(0.0146) |

**Supplementary Table 10.** Precision and recall rates of variants imputed using the TOPMed reference panel inside WES target regions. P - precision. R - recall. For WEGS, this table reports average numbers for each sample. Each sample was sequenced 4 times using WEGS_4P,2X_. HG002 and HG004 were sequenced 5 times using WEGS_8P,5X_. HG003 was sequenced 6 times using WEGS_8P,5X_. The percent of missed true variants is equal to (1 - recall) * 100.

| Sample | TOPMed imputed |  |  |  |  | WEGS 4P, 2X |  |  |  |  | WEGS 8P, 5X |  |  |  |  |
| --- | --- | --- | --- | --- | --- | --- | --- | --- | --- | --- | --- | --- | --- | --- | --- |
|  | TP | FN | FP | P | R | TP (SE) | FN (SE) | FP (SE) | P (SE) | R (SE) | TP (SE) | FN (SE) | FP (SE) | P (SE) | R (SE) |
| SNVs |  |  |  |  |  |  |  |  |  |  |  |  |  |  |  |
| HG002 | 21,458 | 1,285 | 165 | 0.9924 | 0.9435 | 22,379<br>(7) | 364<br>(7) | 446<br>(6) | 0.9805<br>(0.0002) | 0.9840<br>(0.0003) | 22,390<br>(5) | 353<br>(5) | 449<br>(7) | 0.9803<br>(0.0003) | 0.9845<br>(0.0002) |
| HG003 | 21,477 | 1,112 | 111 | 0.9949 | 0.9508 | 22,231<br>(9) | 358<br>(9) | 425<br>(4) | 0.9813<br>(0.0001) | 0.9841<br>(0.0004) | 22,249<br>(6) | 340<br>(6) | 406<br>(5) | 0.9821<br>(0.0002) | 0.9850<br>(0.0003) |
| HG004 | 21,228 | 1,315 | 181 | 0.9915 | 0.9417 | 22,195<br>(9) | 348<br>(9) | 406<br>(7) | 0.9820<br>(0.0003) | 0.9846<br>(0.0004) | 22,197<br>(5) | 346<br>(5) | 402<br>(3) | 0.9822<br>(0.0001) | 0.9847<br>(0.0002) |
| InDels |  |  |  |  |  |  |  |  |  |  |  |  |  |  |  |
| HG002 | 271 | 411 | 8 | 0.9713 | 0.3974 | 648<br>(3) | 34<br>(3) | 57<br>(1) | 0.9197<br>(0.0017) | 0.9498<br>(0.0050) | 647<br>(2) | 35<br>(2) | 58<br>(3) | 0.9177<br>(0.0038) | 0.9481<br>(0.0028) |
| HG003 | 257 | 433 | 8 | 0.9698 | 0.3725 | 649<br>(2) | 41<br>(2) | 52<br>(2) | 0.9254<br>(0.0019) | 0.9409<br>(0.0027) | 651<br>(1) | 39<br>(1) | 52<br>(1) | 0.9265<br>(0.0018) | 0.9432<br>(0.0019) |
| HG004 | 259 | 384 | 7 | 0.9738 | 0.4028 | 616<br>(2) | 28<br>(2) | 43<br>(2) | 0.9357<br>(0.0035) | 0.9572<br>(0.0026) | 615<br>(2) | 28<br>(2) | 40<br>(2) | 0.9393<br>(0.0033) | 0.9568<br>(0.0031) |

**Supplementary Table 11.**
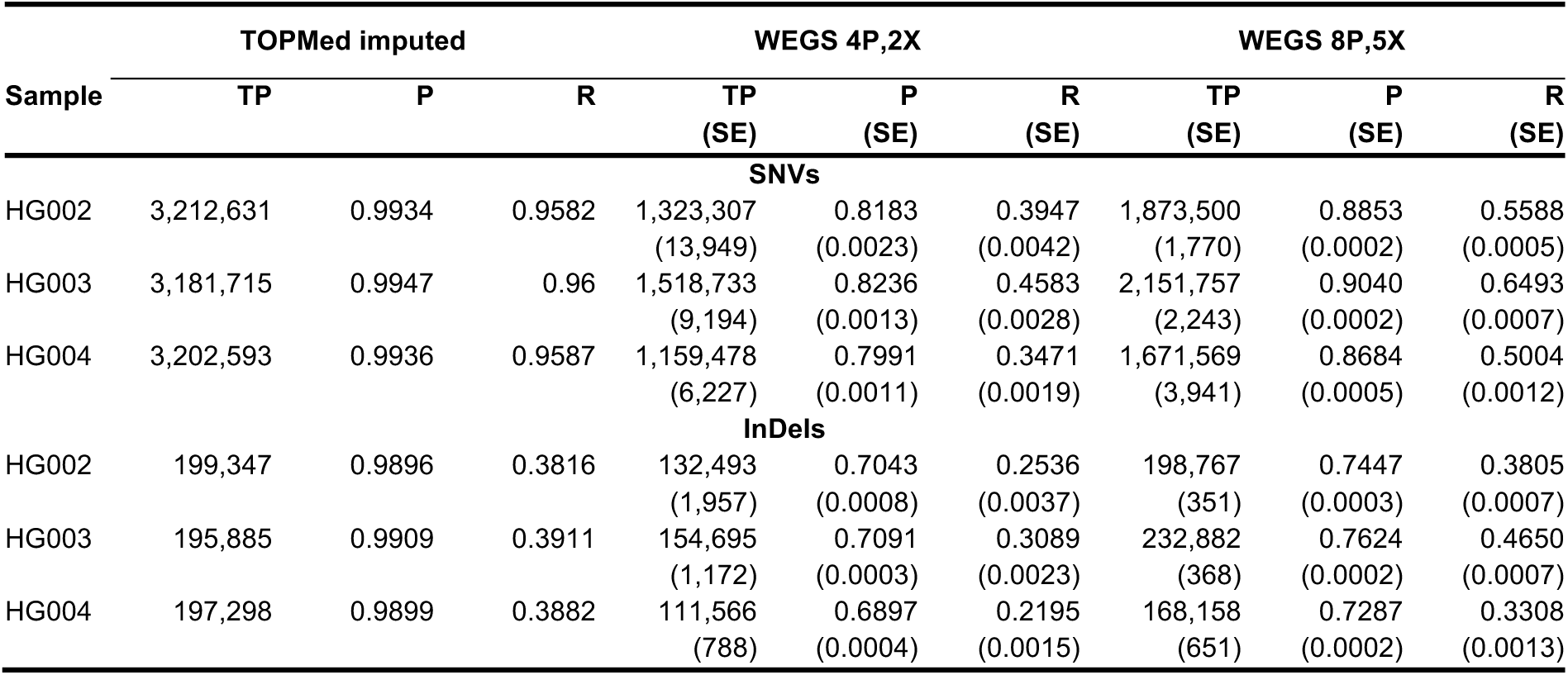
Precision and recall rates of variants imputed using the TOPMed reference panel genome-wide. P - precision. R - recall. For WEGS, this table reports average numbers for each sample. Each sample was sequenced 4 times using WEGS_4P,2X_. HG002 and HG004 were sequenced 5 times using WEGS_8P,5X_. HG003 was sequenced 6 times using WEGS_8P,5X_. The percent of missed true variants is equal to (1 - recall) * 100.

**Supplementary Table 12.**
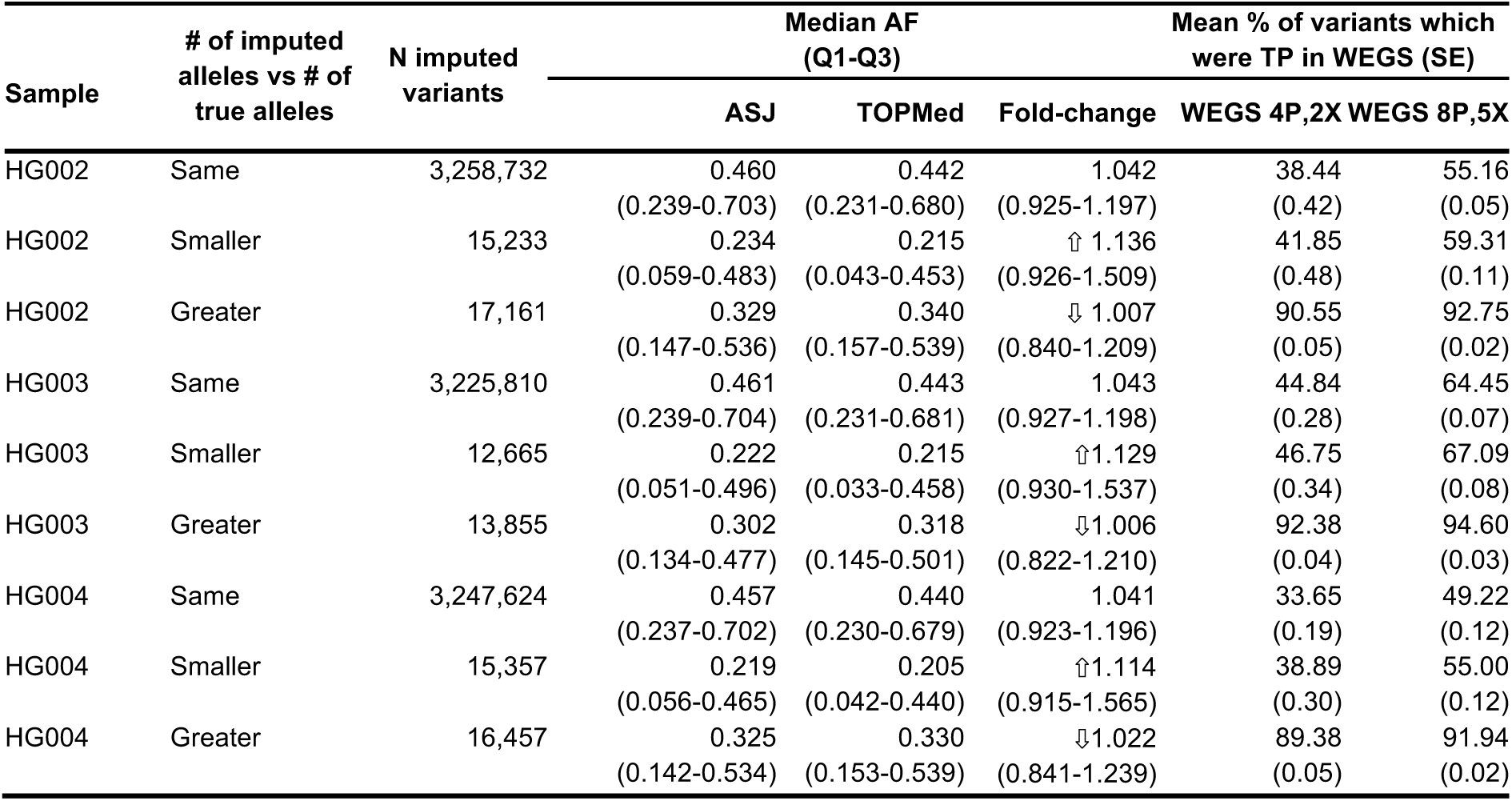
Imputed variants, their allele frequencies, and overlap with true positive (TP) variants in WEGS outside WES target regions. The arrows ⇧ and ⇩ denote the increase and decrease in AF fold-change (AF ASJ / AF TOPMed) compared to variants where the number of imputed alleles matched the number of true alleles.

| Sample | # of imputed alleles vs # of true alleles | N imputed variants | Median AF (Q1-Q3) |  | Fold-change | Mean % of variants which were TP in WEGS (SE) |  |
| --- | --- | --- | --- | --- | --- | --- | --- |
|  |  |  | ASJ | TOPMed |  | WEGS 4P,2X | WEGS 8P,5X |
| HG002 | Same | 3,258,732 | 0.460<br>(0.239-0.703) | 0.442<br>(0.231-0.680) | 1.042<br>(0.925-1.197) | 38.44<br>(0.42) | 55.16<br>(0.05) |
| HG002 | Smaller | 15,233 | 0.234<br>(0.059-0.483) | 0.215<br>(0.043-0.453) | ↑ 1.136<br>(0.926-1.509) | 41.85<br>(0.48) | 59.31<br>(0.11) |
| HG002 | Greater | 17,161 | 0.329<br>(0.147-0.536) | 0.340<br>(0.157-0.539) | ↓ 1.007<br>(0.840-1.209) | 90.55<br>(0.05) | 92.75<br>(0.02) |
| HG003 | Same | 3,225,810 | 0.461<br>(0.239-0.704) | 0.443<br>(0.231-0.681) | 1.043<br>(0.927-1.198) | 44.84<br>(0.28) | 64.45<br>(0.07) |
| HG003 | Smaller | 12,665 | 0.222<br>(0.051-0.496) | 0.215<br>(0.033-0.458) | ↑ 1.129<br>(0.930-1.537) | 46.75<br>(0.34) | 67.09<br>(0.08) |
| HG003 | Greater | 13,855 | 0.302<br>(0.134-0.477) | 0.318<br>(0.145-0.501) | ↓ 1.006<br>(0.822-1.210) | 92.38<br>(0.04) | 94.60<br>(0.03) |
| HG004 | Same | 3,247,624 | 0.457<br>(0.237-0.702) | 0.440<br>(0.230-0.679) | 1.041<br>(0.923-1.196) | 33.65<br>(0.19) | 49.22<br>(0.12) |
| HG004 | Smaller | 15,357 | 0.219<br>(0.056-0.465) | 0.205<br>(0.042-0.440) | ↑ 1.114<br>(0.915-1.565) | 38.89<br>(0.30) | 55.00<br>(0.12) |
| HG004 | Greater | 16,457 | 0.325<br>(0.142-0.534) | 0.330<br>(0.153-0.539) | ↓ 1.022<br>(0.841-1.239) | 89.38<br>(0.05) | 91.94<br>(0.02) |

**Supplementary Table 13.**
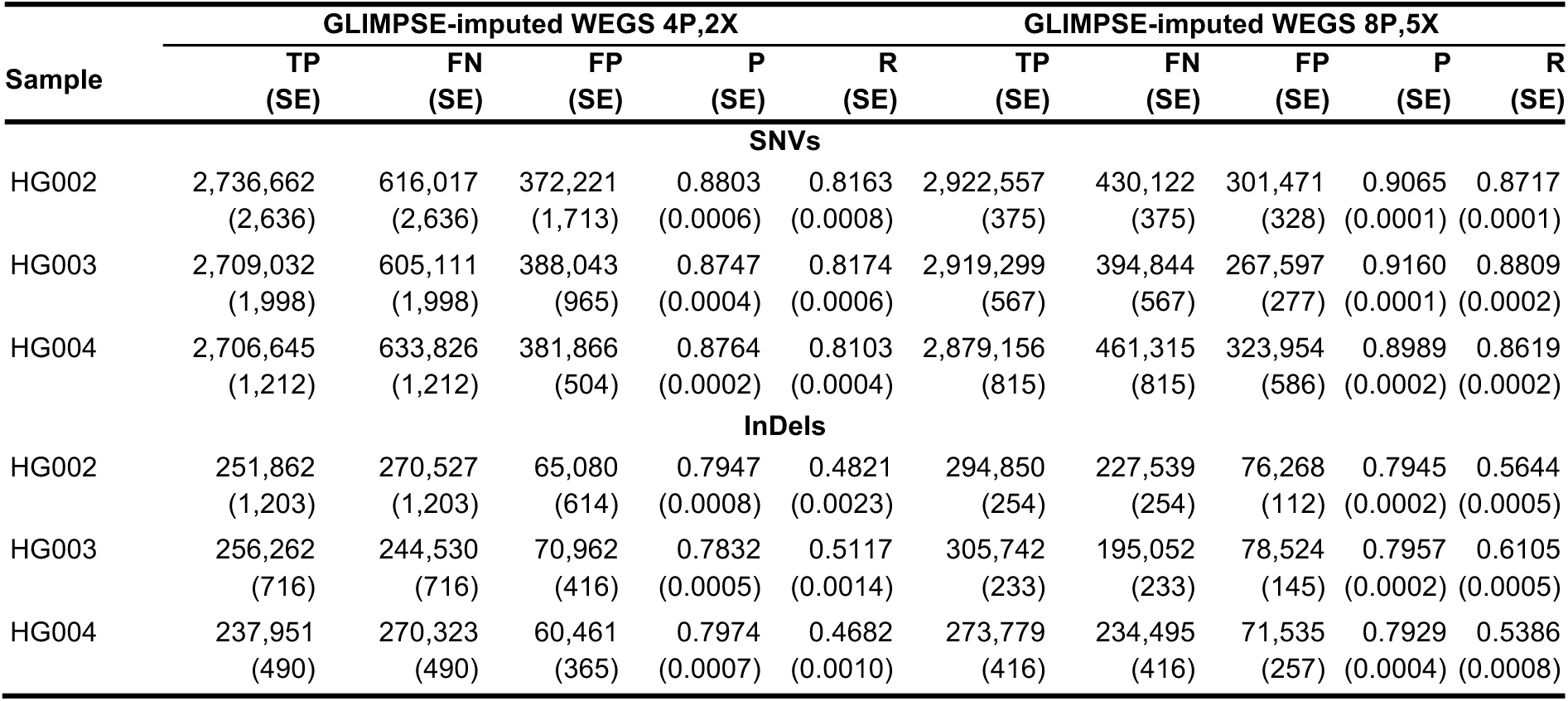
Precision and recall rates of variants imputed using the GLIMPSE method genome-wide. P - precision. R - recall. This table reports the average number for each sample. Each sample was sequenced 4 times using WEGS_4P,2X_. HG002 and HG004 were sequenced 5 times using WEGS_8P,5X_. HG003 was sequenced 6 times using WEGS_8P,5X_. The percent of missed true variants equals (1 - recall) * 100. The local imputation reference panel combines haplotypes from the 1000 Genomes Project and Human Genome Diversity Project (see Methods).

**Supplementary Table 14.**
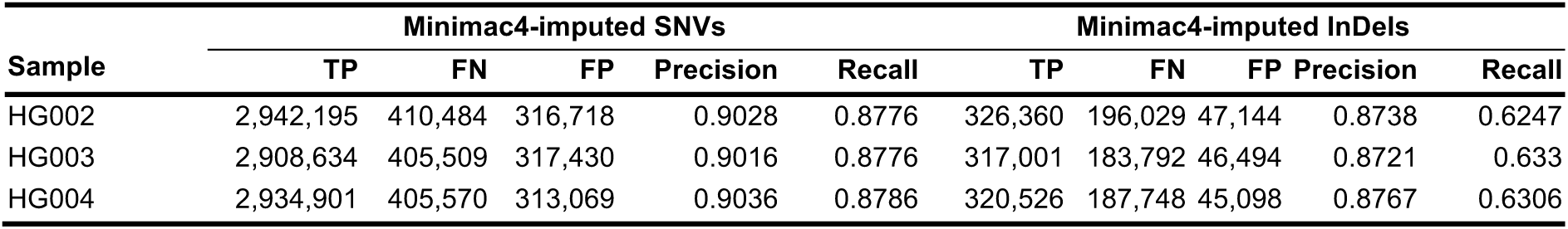
Precision and recall rates of variants imputed using the Minimac4 method genome-wide. The percent of missed true variants equals (1 - recall) * 100. The local imputation reference panel combines haplotypes from the 1000 Genomes Project and Human Genome Diversity Project (see Methods).

| Sample | Minimac4-imputed SNVs |  |  |  |  | Minimac4-imputed InDels |  |  |  |  |
| --- | --- | --- | --- | --- | --- | --- | --- | --- | --- | --- |
|  | TP | FN | FP | Precision | Recall | TP | FN | FP | Precision | Recall |
| HG002 | 2,942,195 | 410,484 | 316,718 | 0.9028 | 0.8776 | 326,360 | 196,029 | 47,144 | 0.8738 | 0.6247 |
| HG003 | 2,908,634 | 405,509 | 317,430 | 0.9016 | 0.8776 | 317,001 | 183,792 | 46,494 | 0.8721 | 0.633 |
| HG004 | 2,934,901 | 405,570 | 313,069 | 0.9036 | 0.8786 | 320,526 | 187,748 | 45,098 | 0.8767 | 0.6306 |

**Supplementary Table 15.**
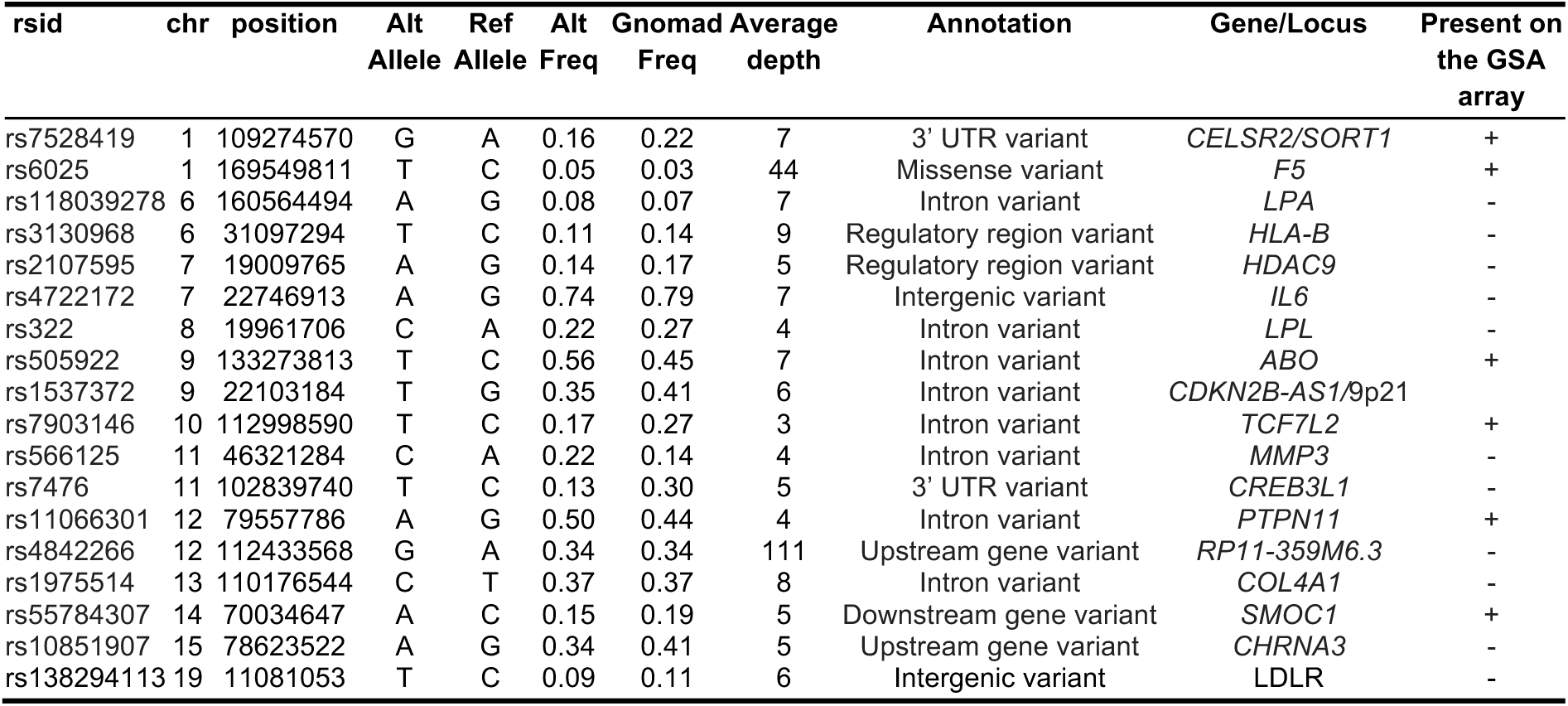
Overview of genome-wide significant loci associated with peripheral artery disease (PAD) in the 862 WEGS sequenced patients. Abbreviations: chr-chromosome; alt-alternative; ref-reference; freq-frequency. GSA-global screening array (24v3). This table reports the allele frequency and average depth of known genome-wide significant peripheral artery disease loci in the WEGS.

**Supplementary Table 16.** Genome-wide significant loci associated with peripheral artery disease (PAD) and the number of variants within the loci present in WEGS and TOPMed. For each locus, we counted the number of variants surrounding the lead variant (rsid) within ±500 kilobase (kb) distance.

| rsid | chr | position | # of SNVs<br>in WEGS | # of SNVs<br>in TOPMed | # of WEGS SNVs absent from TOPMed |  |  |  |  |  |
| --- | --- | --- | --- | --- | --- | --- | --- | --- | --- | --- |
|  |  |  |  |  | Total | Synon | Non-synon | Stop/Splice | Frameshift | Inframe |
| rs7528419 | 1 | 109274570 | 17,631 | 258,859 | 4,481 | 17 | 35 | 2 | 2 | 0 |
| rs6025 | 1 | 169549811 | 14,663 | 254,419 | 3,310 | 9 | 8 | 3 | 0 | 0 |
| rs118039278 | 6 | 160564494 | 15,028 | 272,156 | 3,141 | 1 | 17 | 0 | 0 | 0 |
| rs3130968 | 6 | 31097294 | 32,930 | 255,957 | 6,085 | 24 | 64 | 5 | 10 | 10 |
| rs2107595 | 7 | 19009765 | 14,424 | 322,455 | 2,379 | 4 | 2 | 0 | 0 | 0 |
| rs4722172 | 7 | 22746913 | 15,796 | 276,263 | 3,505 | 3 | 7 | 1 | 3 | 0 |
| rs322 | 8 | 19961706 | 17,333 | 333,151 | 3,251 | 4 | 10 | 1 | 0 | 0 |
| rs505922 | 9 | 133273813 | 14,233 | 290,358 | 4,274 | 9 | 29 | 1 | 1 | 3 |
| rs1537372 | 9 | 22103184 | 20,034 | 324,269 | 2,731 | 3 | 2 | 0 | 0 | 0 |
| rs7903146 | 10 | 112998590 | 14,599 | 268,399 | 3,304 | 1 | 3 | 0 | 0 | 0 |
| rs566125 | 11 | 46321284 | 16,485 | 262,361 | 4,932 | 8 | 34 | 1 | 0 | 0 |
| rs7476 | 11 | 102839740 | 15,023 | 274,797 | 3,198 | 4 | 23 | 0 | 1 | 0 |
| rs11066301 | 12 | 79557786 | 55,094 | 262,291 | 3,143 | 1 | 3 | 0 | 0 | 0 |
| rs4842266 | 12 | 112433568 | 16,233 | 250,632 | 5,016 | 3 | 9 | 2 | 0 | 0 |
| rs1975514 | 13 | 110176544 | 17,779 | 288,888 | 3,274 | 7 | 25 | 1 | 0 | 2 |
| rs55784307 | 14 | 70034647 | 15,766 | 264,425 | 3,387 | 4 | 12 | 0 | 0 | 0 |
| rs10851907 | 15 | 78623522 | 18,510 | 275,484 | 4,514 | 7 | 25 | 1 | 0 | 2 |
| rs62084752 | 17 | 68093252 | 21,373 | 285,583 | 5,903 | 7 | 11 | 0 | 2 | 0 |
| rs138294113 | 19 | 11081053 | 25,994 | 306,166 | 7,227 | 21 | 34 | 3 | 4 | 2 |

